# Phosphoproteomics reveals content and signaling differences between neonatal and adult platelets

**DOI:** 10.1101/2023.09.13.557268

**Authors:** Christopher S Thom, Patricia Davenport, Hossein Fazelinia, Zhi-Jian Liu, Haorui Zhang, Hua Ding, Jennifer Roof, Lynn A Spruce, Harry Ischiropoulos, Martha Sola-Visner

**Author notes:** These authors contributed equally to this work (Christopher S Thom, Patricia Davenport). **Corresponding Authors:** Christopher S Thom, MD, PhD Children’s Hospital of Philadelphia 10-052 Colket Translational Research Building 3501 Civic Center Blvd Philadelphia, PA 19104; Martha Sola-Visner, MD Boston Children’s Hospital Division of Newborn Medicine, Enders Rm. 961 300 Longwood Avenue Boston, MA 02115.

## Abstract

**Background and Objective:** Recent clinical studies have shown that transfusions of adult platelets increase morbidity and mortality in preterm infants. Neonatal platelets are hyporesponsive to agonist stimulation, and emerging evidence suggests developmental differences in platelet immune functions. This study was designed to compare the proteome and phosphoproteome of resting adult and neonatal platelets.

**Methods:** We isolated resting umbilical cord blood-derived platelets from healthy full term neonates (n=9) and resting blood platelets from healthy adults (n=7), and compared protein and phosphoprotein contents using data independent acquisition mass spectrometry.

**Results:** We identified 4745 platelet proteins with high confidence across all samples. Adult and neonatal platelets clustered separately by principal component analysis. Adult platelets were significantly enriched for immunomodulatory proteins, including β2 microglobulin and CXCL12, whereas neonatal platelets were enriched for ribosomal components and proteins involved in metabolic activities. Adult platelets were enriched for phosphorylated GTPase regulatory enzymes and proteins participating in trafficking, which may help prime them for activation and degranulation. Neonatal platelets were enriched for phosphorylated proteins involved in insulin growth factor signaling.

**Conclusions:** Using state-of-the-art mass spectrometry, our findings expanded the known neonatal platelet proteome and identified important differences in protein content and phosphorylation compared with adult platelets. These developmental differences suggested enhanced immune functions for adult platelets and presence of a molecular machinery related to platelet activation. These findings are important to understanding mechanisms underlying key platelet functions as well as the harmful effects of adult platelet transfusions given to preterm infants.

## Introduction

Platelets are anucleate cells generated from megakaryocytes (MKs). Platelets first appear in the fetal circulation at 8-weeks gestation and, by the time of birth, both preterm and term neonates have mean platelet counts within the normal adult range. While platelets are the primary cellular component of hemostasis, over the past two decades it has become increasingly recognized that they also play important roles in angiogenesis, inflammation, and host defense [1-6]. The immune and inflammatory functions of platelets are complex, involving both the innate and adaptive immune systems, which they modulate in a highly context-dependent manner. Platelets express surface and cytoplasmic pattern recognition receptors [7-10]; interact directly with pathogens, immune cells, and complement [1, 11-20]; release antimicrobial molecules as well as pro-inflammatory chemokines and cytokines [21]; and can act as antigen presenting cells [22]. The proteins required to carry out the varied platelet functions come from three main sources - proteins derived from MKs, circulating proteins taken up by the platelets, and proteins actively translated in the platelet itself. Despite the lack of a nucleus, platelets contain thousands of transcripts derived from MKs and the required machinery for protein translation (e.g., rough endoplasmic reticulum, polyribosomes, and other key factors for translation initiation, termination, and regulation) [23-26].

Neonatal platelets (including those from infants born premature) are structurally similar to adult platelets but have key functional differences. Neonatal platelets are hyporesponsive to most platelet agonists, including thromboxane (TXA_2_), ADP, thrombin, collagen, and epinephrine. The mechanisms underlying this hypo-reactivity vary by agonist, but include decreased receptor expression (e.g., PAR1, PAR4, CLEC2, GPVI, and α-adrenergic receptors), decreased GTPase activity, and/or intracellular signaling defects [27, 28]. No prior study has explored whether there are differences in intracellular signaling pathways between resting neonatal and adult platelets that could contribute to the neonatal phenotype.

Within the context of hemostasis, the neonatal platelet hyporeactivity is not due to a developmental deficiency or immaturity. Rather, these properties are integral to a different but uniquely balanced neonatal hemostatic system, termed ‘developmental hemostasis’, in which the neonatal platelet hyporeactivity is counteracted by factors in neonatal blood that enhance platelet-vessel wall interactions, such as increased vWF levels, higher hematocrit, and higher erythrocyte mean corpuscular volume [29]. As a consequence, *in vivo* bleeding times and *in vitro* closure times measured using the Platelet Function Analyzer (PFA-100®) are *shorter* in healthy full term neonates compared to adults [30, 31].

Whether neonatal platelets are part of a similar developmentally-regulated immune system balance is unknown. However, a key developmental difference between neonatal and adult platelets is the differential expression of P-selectin on the platelet surface upon activation. Activated adult platelets translocate more P-selectin to the cell surface compared to neonatal platelets. This makes P-selectin available to interact with its receptor, PSGL-1, which is found on many cells including neutrophils and monocytes [11, 12]. This difference in P-selectin surface exposure suggests that activated adult platelets may be primed for increased immune cell interactions compared to neonatal platelets.

The differences between neonatal and adult platelets are of clinical importance since, when transfused, neonates invariably receive platelets from adult donors. In 2019, the largest randomized trial of platelet transfusion thresholds in preterm neonates (PlaNeT-2) found that neonates randomized to receive a transfusion when the platelet count fell below 50×10^9^/L *had a significantly higher incidence of death and/or major bleeding* compared to neonates randomized to a more restrictive threshold of 25×10^9^/L [32]. The mechanisms mediating these deleterious effects are unknown, but they are likely –at least in part-related to differences between neonatal and adult platelets and the potentially harmful “developmental mismatch” associated with transfusion of adult platelets into neonates. Identifying molecular differences between neonatal and adult platelets has implications for defining the effects of adult platelets on the neonatal hemostatic and immune systems, and could provide insights into why platelet transfusions are harmful.

Two groups previously reported a high concordance in mRNA content when comparing platelets isolated from neonatal umbilical cord blood and adult peripheral blood [33, 34]. However, neonatal platelets were enriched for transcripts associated with protein synthesis, trafficking, and degradation while containing fewer transcripts for genes related to calcium transport, metabolism, actin cytoskeleton reorganization, and cell signaling [33]. In the more recent study, two genes (DEFA3 and HBG1) were identified as platelet biomarkers of neonatal megakaryopoiesis, potentially differentiating neonatal from adult platelets [34]. One prior study also compared the neonatal and adult platelet proteomes and found many proteins essential for platelet function to be equally expressed between neonatal and adult platelets. Examination of differentially expressed proteins showed that neonatal platelets contain fewer proteins related to cellular signaling and increased abundance of proteins related to energy metabolic processes [35].

The current study was designed to expand our understanding of developmental differences in platelet protein content and signaling pathways using state-of-the-art data-independent acquisition (DIA) mass spectrometry, coupled with the first phosphoproteomics study in resting neonatal and adult platelets. Our findings shed light on proteins and signaling pathways that differ between adult and neonatal platelets, and begin to reveal key molecules underlying developmentally regulated differences in platelet function.

## Methods

### Subjects

This study was approved by the institutional review board at Boston Children’s Hospital and at Beth Israel Deaconess Medical Center (BIDMC). Cord blood (CB) was collected from healthy full term infants born by elective cesarean section at BIDMC following an uncomplicated pregnancy. Adult blood was collected from the antecubital vein of healthy adult volunteers who had not taken any anti-platelet medications during the ten days prior to the study. All neonatal and adult blood samples were gently drawn through a 21-gauge needle into 30mL plastic syringes containing 5mL of acid citrate dextrose (ACD) and processed immediately upon collection. One mL of whole blood was set aside for cell counts and for flow cytometric analysis.

### Platelet preparation

Platelets were isolated using the platelet rich plasma (PRP) method. Briefly, collected whole blood was treated with 500nM PGE-1 and 6mM EDTA and centrifuged at 150g for 15 minutes to separate the platelets. The platelet rich plasma (PRP) was removed and centrifuged at 120g for 10 minutes to remove any additional leukocytes and red blood cells. The pure PRP was then centrifuged at 280g for 30 minutes to pellet the platelets. Pelleted platelets were washed twice in PBS containing 2mM EDTA and 100nM PGE-1. The final platelet pellet was flash frozen in liquid nitrogen and then stored at -80°C until processing.

### Platelet activation status and cell purity

Platelet P-selectin surface expression was determined by flow cytometry. Briefly, 10uL of diluted whole blood and final washed platelets were incubated with CD41-APC (BD Bio Sciences 559777) and CD62P-PE (Bio Rad MCA2419PE) antibodies. An IgG-PE control served as the negative control and a sample of whole blood activated with 100ug/mL of thrombin receptor activating peptide (TRAP) served as positive control. P-selectin surface expression was recorded as mean fluorescent intensity (MFI). Platelet purity was determined by incubating 5*μ*L of washed platelets with CD41-APC (BD Bio Sciences 559777) and CD45-FITC (BD Bio Sciences 555482) antibodies to ensure against leukocyte contamination in the final sample.

### Protein extraction

Platelet pellets were prepared in 8M urea lysis buffer as described [36] (**Supplementary Methods**). The protein concentration of each supernatant was assessed by Micro BCA assay (Thermo Scientific).

### Protein hydrolysis

One milligram (1mg) protein from each sample was spiked with 1*μ*g chicken ovalbumin (1:1000), which served as a phosphoprotein internal standard. Proteins were reduced with 5mM dithiothreitol for 45 min, followed by alkylation with 20 mM iodoacetamide for 45 min in the dark. Lysates were diluted with 50 mM Tris-HCl (pH 8.0) to reduce the total urea concentration to 0.8 M. To achieve proteolysis, samples were incubated with LysC (Wako,129-02541) at a 1:100 (enzyme:protein) ratio at 27°C for 2 h, followed by trypsin (Promega, V5111) at a 1:50 (enzyme:protein) ratio at 27°C overnight. Digests were acidified with formic acid (Thermo Scientific, 28905, final concentration 1%) and centrifuged at 20,000g for 10 min. Supernatants were desalted under a vacuum manifold using an Oasis HLB 96 well plate (60mg, Waters, 186000679) that had been preconditioned with 1 × 400*μ*L 100% acetonitrile and equilibrated with 2 × 400*μ*L 0.1% trifluoroacetic acid (J T Baker, 9470-00). Tryptic peptides were loaded into the plate, washed with 3 x 400*μ*L 0.1% trifluoroacetic acid, and eluted into a protein Lobind plate 96/2000 uL (Eppendorf, EP0030504305) with 3 x 300 *μ*L 50% acetonitrile/0.1% trifluoroacetic acid.

### Allocation of digests for further processing

Five percent (5%) of each sample was used for whole proteome analysis. 75% of each sample was used for IMAC phosphopeptide enrichment [36-38]. The remaining 20% for each sample was pooled and used for phosphopeptide spectral library generation. All peptides were lyophilized and stored at -80°C. Peptides were sub-fractionated by HPLC and subjected to IMAC enrichment and lyophilized [36, 37] (**Supplementary Methods**). Prior to LC-MS/MS analysis, all peptides were solubilized in 0.1% TFA containing iRT peptides (Biognosys AG, iRT).

### Mass spectrometry data acquisition

Samples were analyzed on an Exploris 480 mass spectrometer (Thermo) coupled with an Ultimate 3000 nano UPLC system and an EasySpray source using data independent acquisition settings (**Supplementary Methods**).

### QA/QC and system suitability

The suitability of the Exploris 480 instrument was monitored using QuiC software (Biognosys) for the analysis of the spiked-in iRT peptides (**Supplementary Methods**). These data were analyzed in MaxQuant [39] and the output was subsequently visualized using the PTXQC [39] package to track the quality of the instrumentation.

### Mass spectrometry raw data processing

DIA MS/MS raw files were processed in Spectronaut 16 (Biognosys AG) [40]. We used a reference human proteome containing 20,385 canonical and reviewed isoforms from Uniprot appended with the list of 245 common protein contaminants. Trypsin was specified as the enzyme, with up to two possible missed cleavages. Carbamidomethyl of cysteine was specified as a fixed modification and protein N-terminal acetylation and oxidation of methionine were considered as variable modifications. For the phosphoproteome data, we added phosphorylation of Serine, Threonine, and Tyrosine as variable modifications. A false discovery rate <1% was set for peptide and protein identification. All other search parameters were set to default values. The phosphopeptide spectral library was also generated in Spectronaut using default settings.

### Whole proteome data analysis

We used publicly available R software for data processing and statistical analyses. The MS2 intensity values generated by Spectronaut were used to analyze the whole proteome data. The data were log2-transformed and normalized by subtracting the median for each sample. We filtered the data to have a complete value for a protein in at least one cohort. To compare proteomics data between groups, the limma t-test was employed to identify differentially expressed proteins, and volcano plots were generated in visualize the affected proteins while comparing different groups. Lists of differentially abundant proteins was then sorted based on the adj.P.Value <0.05 and |FC| > 1.5, yielding a prioritized list for downstream bioinformatics analysis.

### Phosphoproteome data analysis

The peptide quantification report was collapsed into modification-specific peptide-like tables using default Spectronaut settings. Using the reported PTMLocalizationProbabilities, the localization cutoff of 0.75 was used to retain the high-confidence sites. Phosphorylation of Ser, Thr, and Tyr, acetylation of protein N-terminal, and oxidation of Met were considered as variable modifications. Phosphopeptide intensities were log2-transformed and normalized to the median for each sample. The limma t-test was used for differential expression analysis of phosphopeptides using adj.P.Value < 0.05 and |FC| > 1.5 as cutoff thresholds. Phosphoprotein quantification was performed by calculating the absolute intensity of the detected phosphopeptides for each protein, and reporting the maximum value as the overall intensity. The phosphoproteins selected by this workflow were carried forward for further analyses.

### Statistical analysis and data plotting

High-confidence proteins and phosphoproteins were analyzed using the R computational environment and established packages (**Table S1**). Pathway enrichment analyses referenced standard Gene Ontology pathways [41]. Related plots were constructed to capture the most enriched pathways. A comprehensive catalogue of pathway enrichment results is presented in the related Supplemental Tables.

Cytoscape (v3.9.1) analysis and plots were created using the Omics Visualizer application [42]. A catalogue of specific phosphopeptides for each protein shown in the interaction plots can be found the Supplement for this manuscript. KEA analysis was performed using the online interface (https://maayanlab.cloud/kea3/). Other individual plots and statistical analyses used R and GraphPad Prism 9. Figures were made using BioRender (www.biorender.org).

### Public data set analysis

We conducted a post hoc analysis of publicly available neonatal and adult platelet proteomic data using the raw data processing, whole proteome data analysis, and data plotting methods described above (http://www.ebi.ac.uk/pride/archive/projects/PXD004578) [35]. To compare proteins identified in this prior analysis vs our study, we queried protein gene names from our results against those retrieved from the previously published neonatal or adult platelet proteomes, and manually created a Venn diagram for comparison.

### Data and code availability

All coding scripts utilized publicly available software packages and are available by request. Mass spectrometry data from this study have been deposited to the ProteomeXchange Consortium via the PRIDE partner repository with the dataset identifier PXD043330.

## Results

### Platelet isolation

We isolated platelets from umbilical cord blood obtained from full term neonates (n=9) and blood obtained from healthy adult volunteers (n=7) (**Figure 1A**). Demographic characteristics of the neonates and adults in the study are provided in **Figure 1B**. No leukocyte contamination (CD45^+^ cells) was found by flow cytometry in the washed platelet samples. We also confirmed the resting state of isolated platelets by P-selectin staining, finding a small increase in P-selectin surface expression following platelet isolation and washing compared to fresh whole blood samples (**Figure 1C-D**). One adult sample exhibited high P-selectin mean fluorescence intensity (MFI) immediately after collection, which persisted after washing (**Figure 1D**), and two neonatal samples had increased platelet P-selectin MFI after washing (**Figure S1A**). However, these samples generally clustered with other developmentally matched samples by principal component analyses (**Figure S1B-C**) and thus were not excluded from this study.

**Figure 1.**
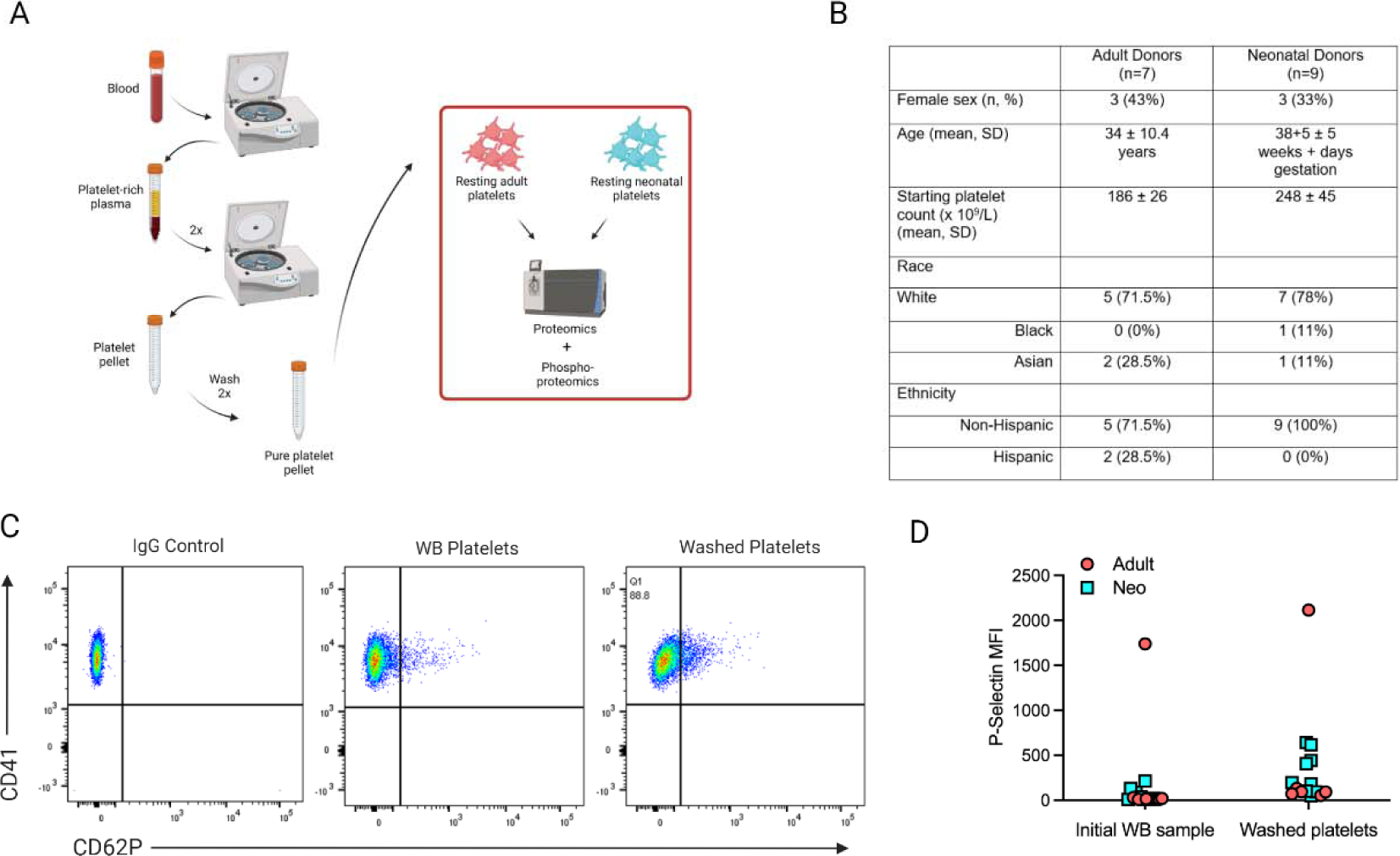
Collection and isolation of resting neonatal and adult platelets. **A**. Schematic overview of sample isolation. **B**. Demographics for neonatal and adult platelet donors for the study. **C**. Representative FACS plot for activation status of platelets in whole blood (WB platelets) prior to washing and isolated platelets following all washing steps (Washed platelets). **D**. P-selectin mean fluorescence intensity (MFI) of platelets in whole blood immediately after collection and following isolation and washing.

### Proteomic analysis reveals differences in immunity, inflammation, metabolism and ribosomal content between resting neonatal vs adult platelets

We obtained the proteome of resting platelet samples by data independent acquisition (DIA) mass spectrometry. We identified 4182 total proteins, a number that is similar to previous estimates of the comprehensive platelet proteome [43] (**Figure S2A-C** and **Table S2**). Each of the 4182 proteins was present in all samples, making this a relatively conservative estimate.

Pathway enrichment analysis of the complete platelet proteome revealed biology critical to platelet function, which was consistent with prior studies [35, 41, 44] (**Figure S3** and **Table S3**). We wanted to compare our results to those of a prior proteomic study of adult and neonatal platelet proteins [35]. When we used consistent methods to analyze our data and prior study results, we found that our results captured 92% of this prior catalogue of adult and neonatal platelet proteins (**Methods**), and that our study expanded the total number of identified platelet proteins by ∼45% (**Figure S4**).

While all platelets were similar in overall protein content, neonatal and adult samples clustered separately on principal component analysis (**Figure 2A**). There were 331 proteins with abundance significantly different between neonatal and adult platelets (fold change*≥*1.5, p<0.05, **Figure 2B-C** and **Table 1**). Pathway analysis of the 169 proteins significantly more abundant in the adult platelet proteome revealed enrichment for immune and inflammatory pathways, as well as proteins found on the cell membrane and those related to signaling and calcium mobilization (**Figure 2D**, **Table S2**, and **Table S4**). Conversely, pathway analysis of the 162 proteins significantly more abundant in neonatal platelets showed enrichment for metabolic pathways and ribosomal components (**Figure 2E**, **Table S2**, and **Table S5**). Specific proteins involved in immune and acute humoral responses that were significantly more abundant in adult compared to neonatal platelets included β-2 microglobulin (β2M), the chemokine CXCL-12 (SDF-1), PDGFA, PDGFB, and TGFβ2 (**Figure 3A-D**). Most complement proteins and related molecules were also enriched on adult platelets (**Figure 3E** and **Figure S5**). Excluding potential outliers (**Figure S1**) from these analyses did not change the results. Interestingly, P-selectin (*SELP*) was not significantly enriched at a total protein level in adult compared to neonatal platelets (1.1-fold, p_adj_=0.2) (**Figure 3A** and **Table S2**).

**Figure 2.**
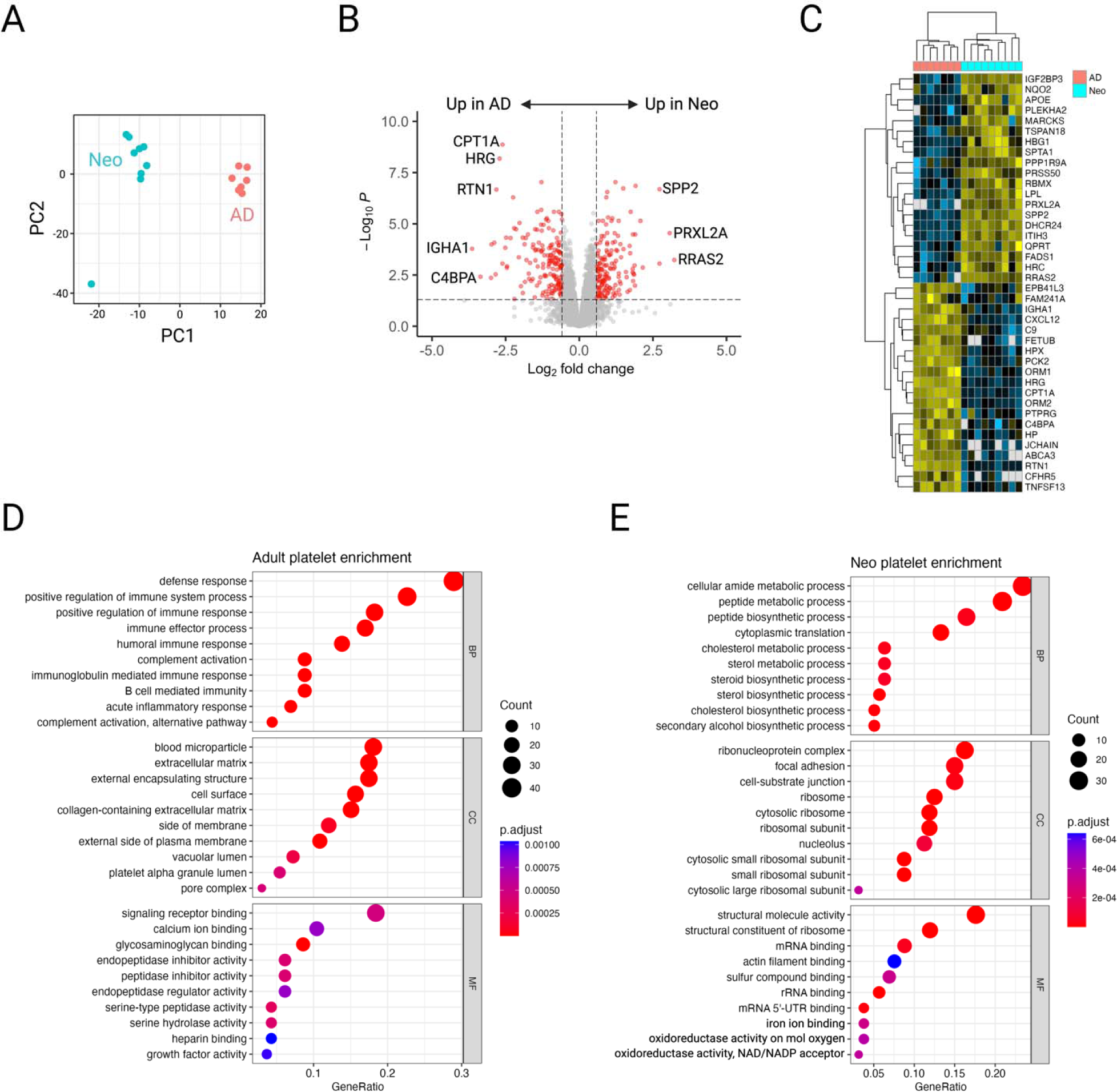
Differences between the neonatal and adult platelet proteomes. **A**. Principal component analysis (PCA) plot for neonatal and adult platelet samples. **B.** Volcano plot depicting statistical significance and changes in protein abundance for neonatal and adult platelets. Proteins with a log (fold-change) >1.5 that met statistical significance (p<0.05) are highlighted in red. Select significantly different proteins are labeled. **C.** Heatmap of the top proteins enriched in adult or neonatal platelets. **D.** Gene ontology pathway analysis for proteins enriched in adult platelets. Significantly enriched biological process (BP), cell compartment (CC), and molecular function (MF)-related pathways are shown. **E.** Gene ontology pathway analysis for proteins enriched in neonatal platelets. Significantly enriched biological process (BP), cell compartment (CC), and molecular function (MF)-related pathways are shown.

**Figure 3.**
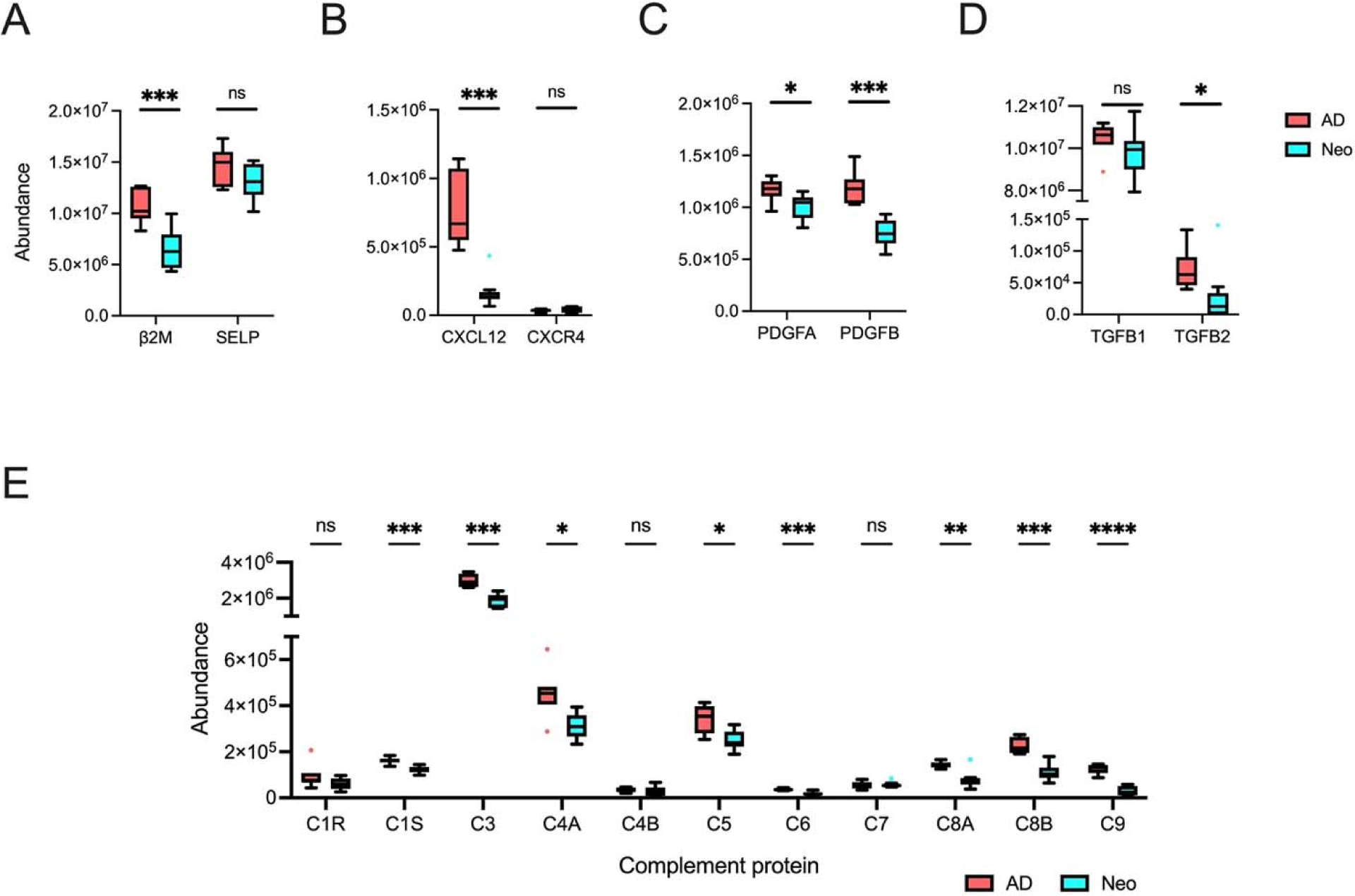
Immunomodulatory and complement proteins are enriched in adult (AD) vs neonatal (Neo) platelets. **A-D**. Comparisons of total protein abundance for **A.** Beta 2-microglobulin (B2M), P-selectin (*SELP*), **B**. CXCL12 and cognate receptor CXCR4, **C.** Platelet derived growth factors A and B, and **D.** Transforming growth factor beta 1 and 2. **E.** Comparisons of total protein abundance for the indicated complement proteins. Tukey box plots depict median with 25^th^-75^th^ interquartile range (IQR), and whiskers represent 1.5 times IQR. Statistical significance was calculated using the Holm step-down procedure. ns, not significant. *p<0.05, **p<0.01, ***p<0.001, ****p<0.0001.

**Table 1.**
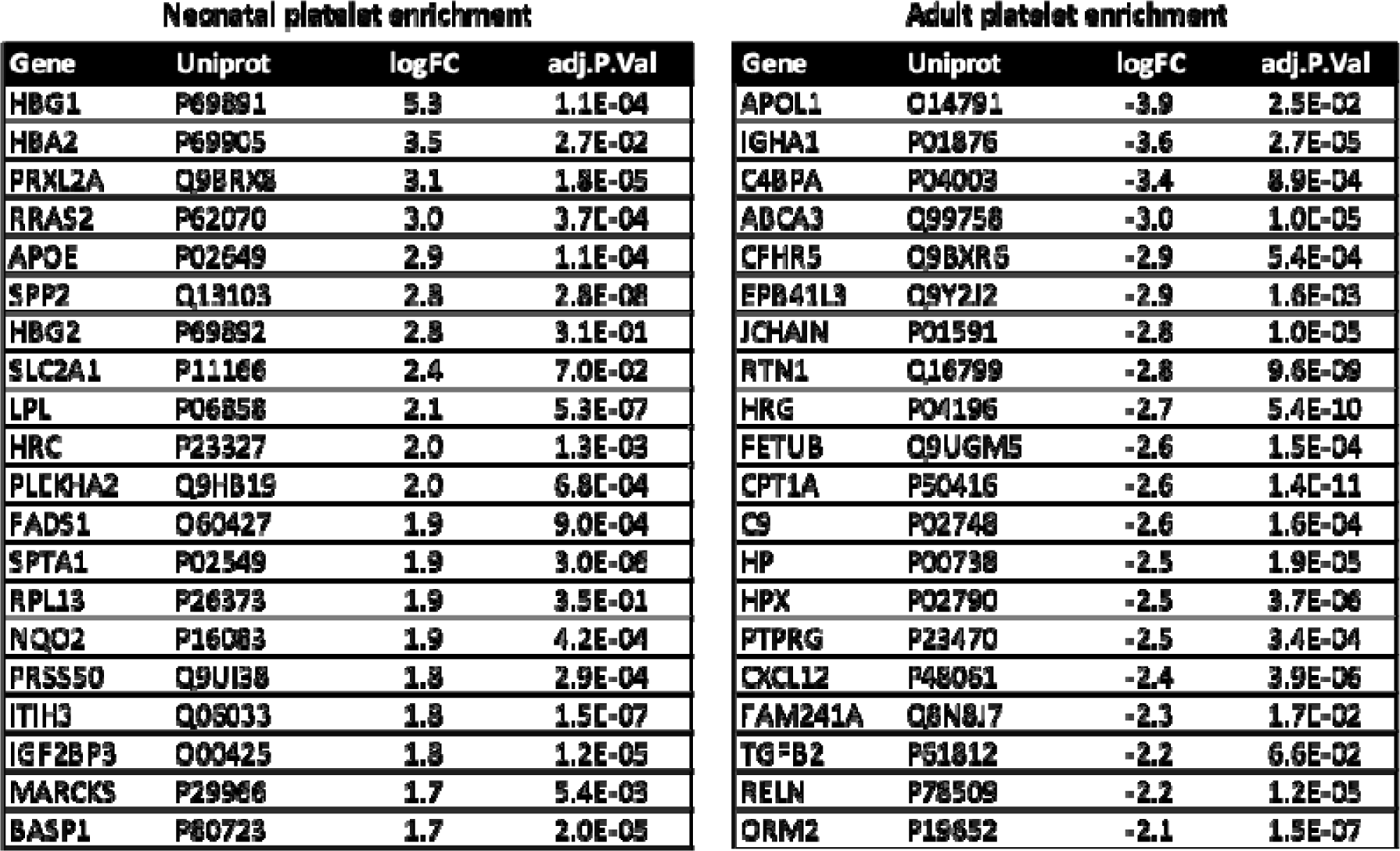
Top 20 most enriched proteins in neonatal and adult platelets. The logFC represents the log_2_(Fold Change) comparing neonatal vs adult platelets.

### Overview of the platelet phosphoproteome

We also wanted to gain insight into molecular signaling processes in neonatal and adult platelets. Protein phosphorylation is the most common post-translational protein modification and at least 30% of all cellular proteins are estimated to be phosphorylated at any given time [45, 46]. We used established pipelines to enrich and detect phosphorylated peptides from neonatal and adult platelet lysates using DIA mass spectrometry [37]. We identified a total of 17,852 phosphopeptides from 2115 phosphorylated proteins across all samples (**Figure 4**, **Figure S6** and **Table S6**). We combined the list of detected phosphoproteins with our initial proteomics results. This combined list of included 4745 individual proteins, 45% of which were phosphorylated (2115/4745).

**Figure 4.**
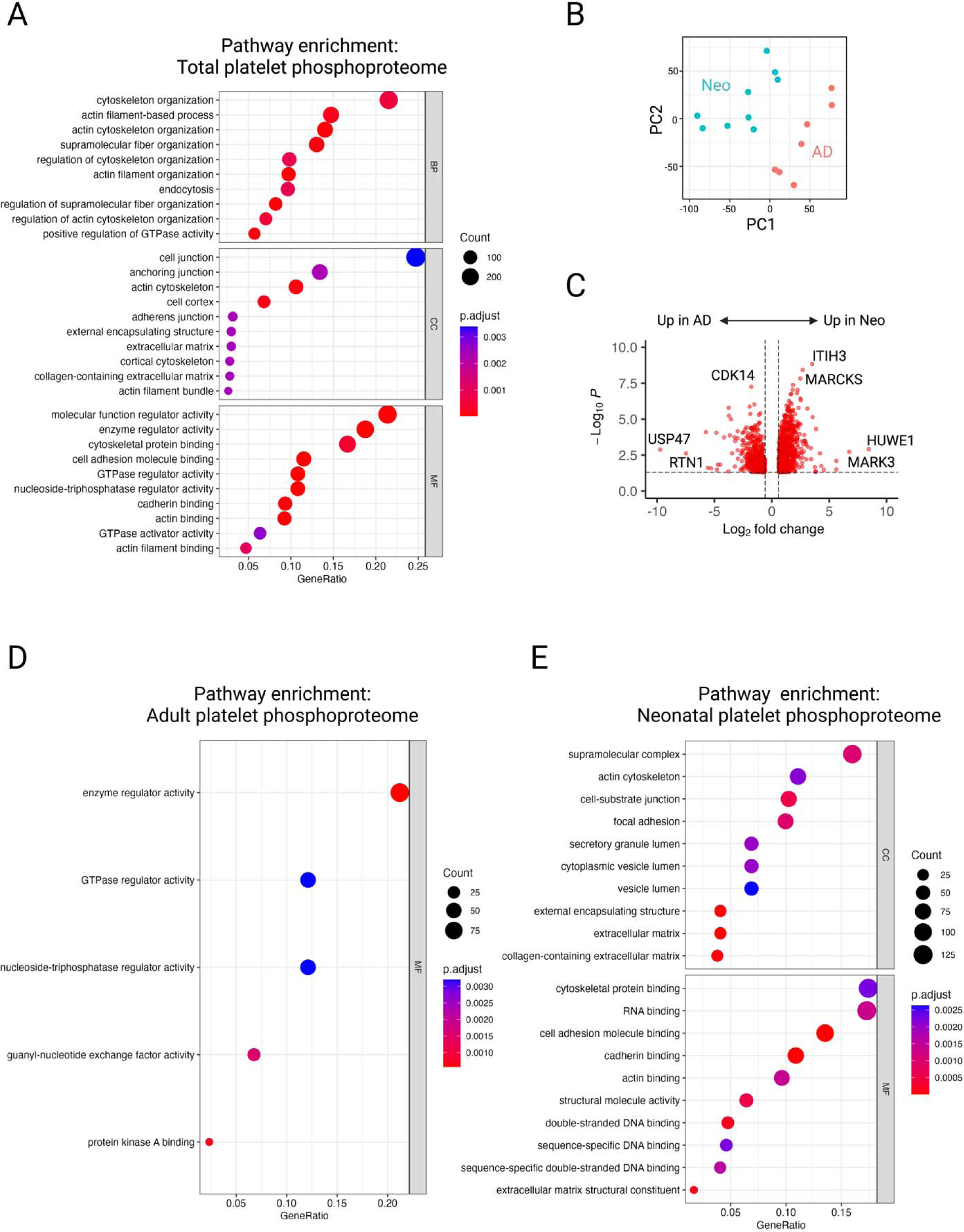
The platelet phosphoproteome is enriched for cytoskeletal and signaling molecules, which differ between neonatal and adult platelets. **A.** Gene Ontology (GO) pathway analysis for phosphoproteins enrichment, considering all phosphopeptides enriched with a fold-change >1.5 that met statistical significance (p<0.05). **B.** Principal component (PC) analysis plot for neonatal and adult platelet samples. **C.** Volcano plot depicting statistical significance and changes in phosphoprotein abundance for neonatal and adult platelets. **D.** Gene ontology pathway analysis for phosphoproteins enriched in adult platelets. Significantly enriched molecular function (MF)-related pathways are shown. There were no significantly enriched biological process (BP) or cell compartment (CC) pathways. **E.** Gene ontology pathway analysis for phosphoproteins enriched in neonatal platelets. Significantly enriched cell compartment (CC) and molecular function (MF)-related pathways are shown. There were no significantly enriched biological process (BP) pathways.

Pathway analysis of the global platelet phosphoproteome revealed enrichment in actin cytoskeletal regulation, cell-cell interactions, and GTPase regulation, among other processes (**Figure 4A** and **Table S7**). These were somewhat different than the pathways enriched in our proteomic analysis (**Figure 2D-E** and **Figure S3**). This likely reflects the importance of phosphorylation cascades in regulating proteins involved in processes such as platelet activation, granule trafficking, and/or degranulation through the identified pathways [47], which may be different from proteins not subjected to phosphorylation (e.g., granule contents).

### Comparison of the neonatal and adult platelet phosphoproteomes reveals differences in GTPase, actin cytoskeletal, and membrane component regulation

Phosphoproteomic analysis showed that neonatal and adult samples clustered separately (**Figure 4B**), with a total of 1183 phosphoproteins found in significantly different abundance between neonatal vs adult samples (fold change*≥*1.5, p<0.05, **Figure 4C**). Pathway analysis of the 445 phosphoproteins significantly more abundant in adult platelets showed enrichment in enzyme regulation, particularly of GTPase enzymes (**Figure 4D** and **Table S8**). The 738 phosphoproteins found in relative higher abundance in neonatal platelets were enriched for cytosplasmic vesicle and membrane localization, including regulatory molecules for focal adhesion, cytoskeletal interactions, and secretory granules (**Figure 4E** and **Table S9**).

### Enriched phosphopeptides in neonatal vs adult platelets suggest proteins and mechanisms underlying differential reactivity and granule secretion

Of the 17,852 phosphopeptides identified in our analysis, 8 phosphopeptides were detected only in neonatal samples but not in *any* adult sample. Conversely, two phosphopeptides were detected only in adult platelets (**Figure 5A**). We reasoned that these were developmental stage-specific phosphopeptides that might reflect key differences between neonatal and adult platelet signaling mechanisms. In support of this, 3 exclusive neonatal phosphopeptides were related to insulin growth factor signaling (e.g., IGF1R, IGF2BP1, IGF2BP3). The RAP1GAP2 phosphopeptide detected in adult platelets may reflect differential Rap signaling activities, known to impact platelet reactivity (**Figure 5A**).

**Figure 5.**
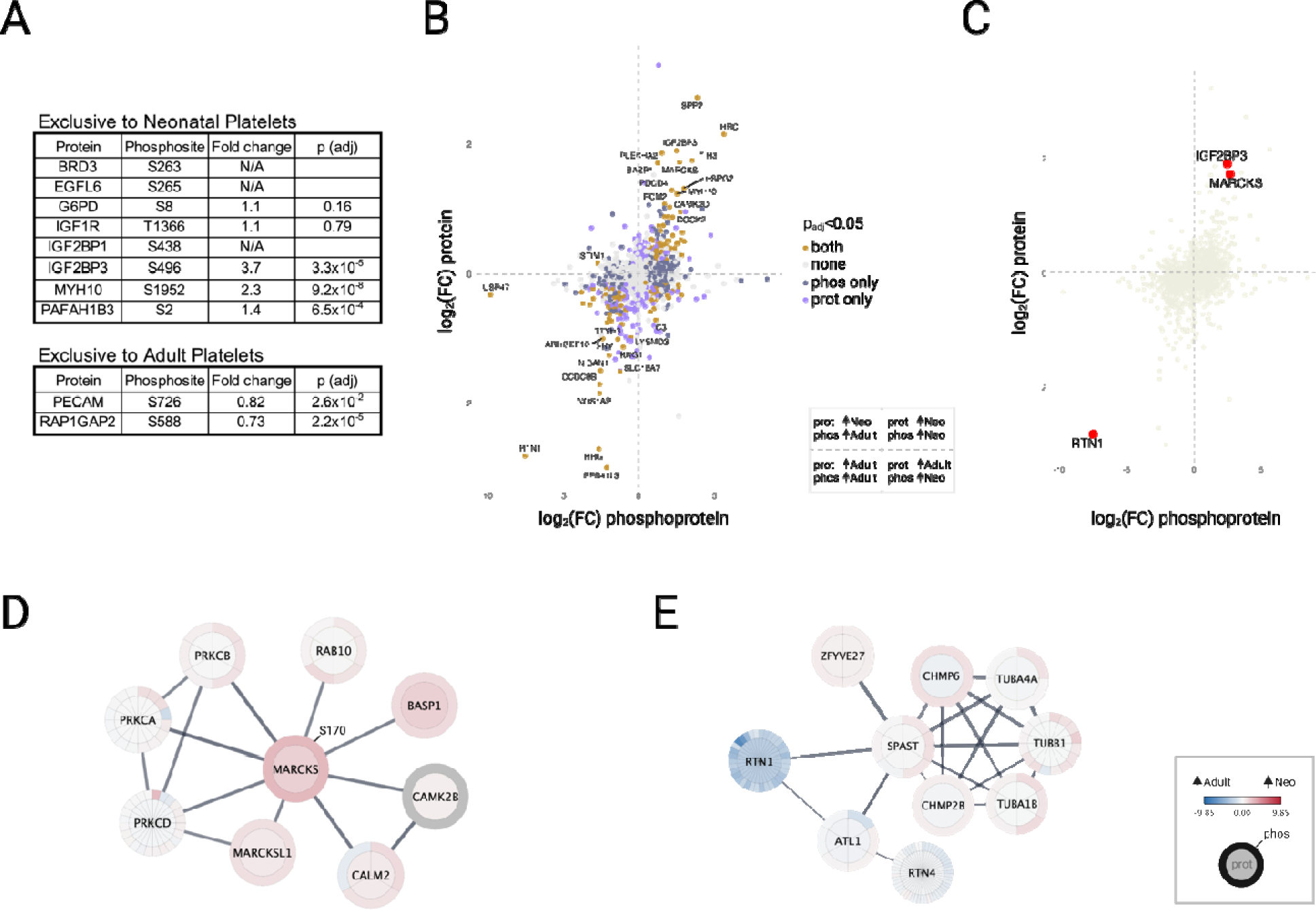
Differential pathway biology revealed by phosphoproteomic analysis of neonatal and adult platelets. **A**. Phosphopeptides that were uniquely present in all neonatal samples but no adult samples, or in all adult samples but no neonatal samples. **B**. Scatterplot depicting relative protein and phosphoprotein abundance in neonatal and adult platelets. Dot colors indicate proteins that met statistical significance (p<0.05) in our proteomic analysis (purple), phosphoproteomic analysis (slate), both (brown), or neither (gray). **C.** Scatterplot highlighting selected proteins (in red) discussed in the text. **D-E.** Cytoscape plots depicting protein interactions for (**D**) MARCKS or (**E**) RTN1 in platelets. Inner circle color depicts total protein enrichment in neonatal vs adult platelets. Outer rim depicts enrichment of individual phosphopeptides in neonatal vs adult platelets.

Alternatively, detection of developmental stage-specific phosphopeptides could reflect the limits of detection in our mass spectrometric analysis, with phosphopeptides more likely to be detected in samples containing higher concentrations of those proteins. In support of this possibility, several exclusive phosphopeptides came from proteins that were significantly enriched in neonatal or adult platelets, and a line of best fit had a significantly positive slope when we plotted protein abundance vs phosphoprotein abundance on a scatterplot (Pr(>|t|) = 2.7×10^-51^, **Figure 5B** and **Figure S7A**). Finally, developmentally regulated protein isoforms could also be responsible for phospho-sites being uniquely present in adult or neonatal samples.

Further analysis of enriched phosphopeptides brought several interesting proteins to our attention, including phosphoproteins that may help explain differences in neonatal and platelet functions (**Figure 5C** and **Figure S7**). Myristolated alanine-rich C kinase substrate (MARCKS) protein (3.3-fold, p_adj_=0.02) and phosphoprotein (6.5-fold, p_adj_=9.5×10^-6^) were significantly enriched in our neonatal platelet samples (**Figure 5C-D**). In contrast, we identified 34 phosphorylated Reticulon 1 (RTN1) peptides, 31 of which were significantly more abundant in adult platelets (each with p<0.05, **Figure 5C-E**). RTN1 isoforms regulate membrane trafficking and interact with SNARE complex proteins [48] (**Figure 5E**). SNARE complex proteins mediate platelet degranulation and were previously implicated in the hyporeactive degranulation phenotype of neonatal platelets [49]. Consistent with these prior findings, SNARE complex proteins were less abundant in neonatal platelets (**Figure S7C**).

### Differential kinase activities in neonatal and adult platelets

We next sought to identify putative kinases responsible for phosphorylation activities in platelets. Kinase enrichment analysis (KEA3) integrates multiple data sources to estimate and rank kinase enzyme activities, allowing robust predictions [50]. Analysis of a combined neonatal and adult platelet phosphoproteome revealed the expected kinases that have been functionally validated in platelets, including the SRC-family kinases FYN, LYN, SRC, and SYK [51] (**Figure 6A**). Cross-regulatory roles among these and other active kinases suggested a complex web of phosphorylation signaling and regulatory activities in our platelet samples (**Figure 6B**).

**Figure 6.**
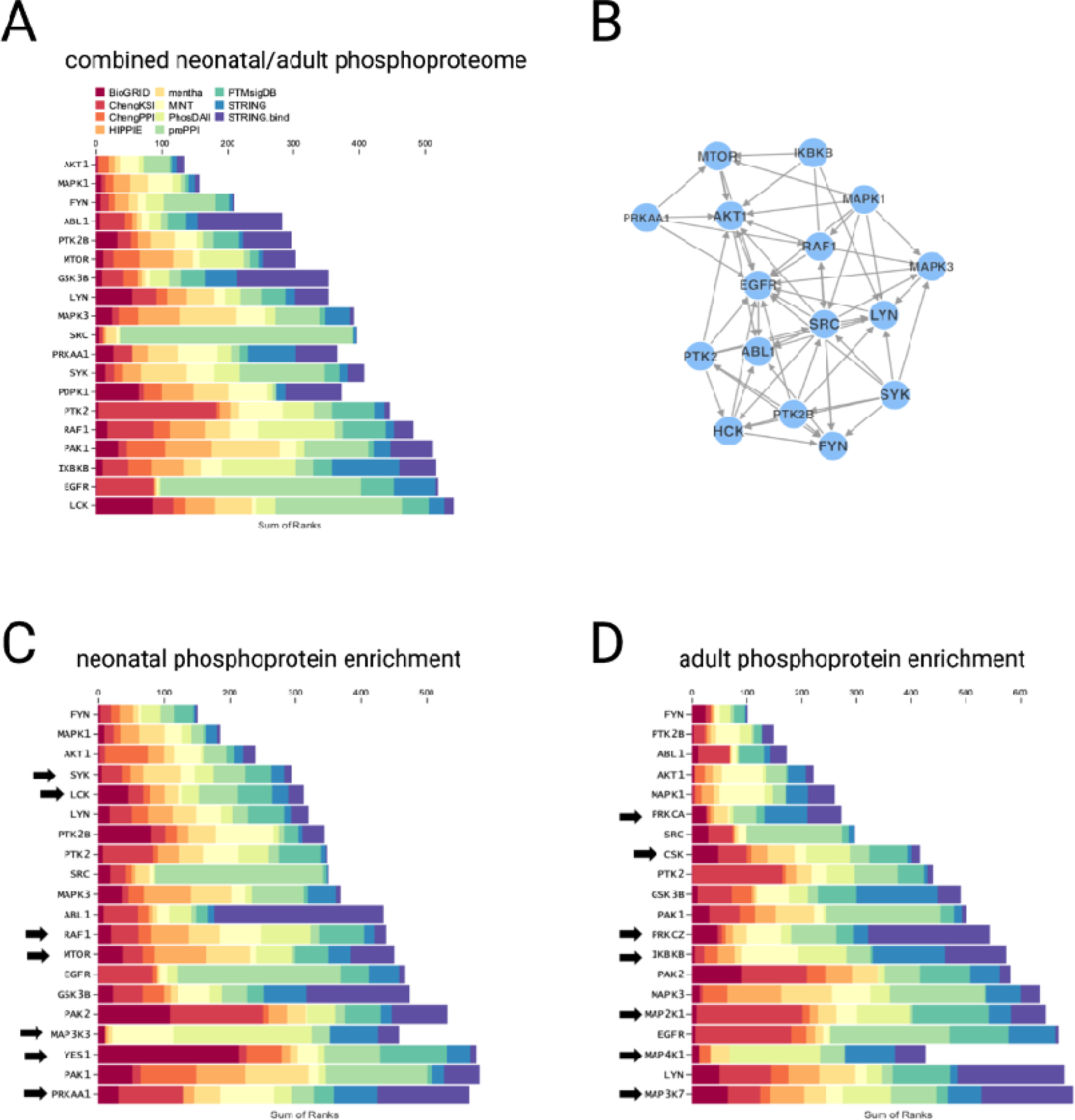
Kinase enrichment analysis underlying activities in the platelet phosphoproteome. **A.** Kinase enrichment analysis for phosphoproteins in the combined neonatal and adult phosphoproteome. Colors represent prioritized ranks from the algorithms depicted at top, with a shorter bar representing a more significant Sum of Ranks from these algorithms. **B.** Cytoscape interaction plot for active platelet kinases as indicated by kinase enrichment analysis. **C.** Active kinases in neonatal platelets based on significantly enriched phosphoproteins. Arrows depict kinases identified among the top 20 most active kinases in neonatal platelets but not adult platelets. **D.** Active kinases in adult platelets based on significantly enriched phosphoproteins. Arrows depict kinases identified among the top 20 most active kinases in adult platelets but not neonatal platelets.

We then focused on kinases inferred to have altered activities from differentially abundant phosphoproteins in neonatal vs adult platelets. Analysis of phosphopeptides significantly more abundant in neonatal platelets identified mTOR and LCK as putatively responsible kinases, among others (**Figure 6C**). Interestingly, mTOR activity coordinates metabolic signaling that we and others have shown to be enriched in neonatal platelets [34] (**Figure 2E**), and we previously found increased mTOR activation in neonatal compared to adult megakaryocytes [52]. LCK is a lymphocyte-specific kinase that we did not detect at the protein level in our study. It is possible that its phosphorylation targets overlap those of other SRC kinases, or that this finding relates to intercellular signaling from interactions between neonatal platelets and circulating lymphocytes [53, 54].

Kinase enrichment also suggested that phosphoproteins enriched in adult platelets were driven in part by activities of several generalized MAP-kinases (**Figure 6D**), resulting in stronger phospho-regulatory effects on GTPase enzymes, actin cytoskeletal components, and membrane regulation pathways revealed by our pathway analyses (**Figure 4D**). An increase in Inhibitor of nuclear factor kappa B kinase subunit beta (IKBKB) activities may be inferred by enhanced immune and inflammatory phosphoproteins, although IKBKB itself was not identified in our platelet proteome or phosphoproteome (**Figure 6D**, **Table S2**, and **Table S6**).

## Discussion

We applied state-of-the-art data independent acquisition mass spectrometry, coupled with phosphoproteomics, to provide a comprehensive characterization of differences in protein content and signaling pathways between resting neonatal and adult platelets. The data increased our understanding of developmental differences in platelet function, having captured 92% of a previously published catalogue of proteins present in neonatal and adult platelets [35] and substantially expanding the total number of identified proteins.

Among the top 20 proteins that were significantly more abundant in neonatal platelets, several have been reported in previous studies comparing neonatal and adult megakaryocytes and/or platelets. These included the gamma globin subunits HBG1 and HBG2 (**Table 1**), which are components of fetal hemoglobin. Previous proteomics [35] and transcriptomics [33, 34] studies also identified erythroid markers in neonatal platelets as well as in cord blood-derived megakaryocytes [55], suggesting incomplete separation between the erythropoietic and megakaryopoietic lineages in fetal and neonatal life. We also found IGF2BP3 to be significantly more abundant in neonatal vs adult platelets, similar to prior reports in neonatal megakaryocytes and consistent with its role as a master regulator of the neonatal megakaryocyte program [55].

Pathway analysis of the 169 proteins that were significantly more abundant in adult platelets revealed an enrichment for immune and inflammatory pathways. Recent studies comparing the mRNA expression profiles of neonatal and adult *murine* megakaryocytes and platelets also found that adult cells were enriched for immune-related pathways [56, 57]. These combined observations strongly suggest that platelets may have different functional specifications at different developmental stages in both mice and humans, with adult platelets having increased immunological functions compared to neonatal platelets. Specific proteins involved in immune and acute inflammatory/humoral responses that were significantly more abundant in adult compared to neonatal platelets included β-2 microglobulin (β2M), PDGFA/B, the chemokine CXCL-12 (SDF-1), and several complement components (**Figure 3**). β2M, a molecular chaperone for the major histocompatibility class I (MHC I) complex, is released upon platelet activation and is a major mediator of monocyte pro-inflammatory differentiation through non-canonical TGFβ receptor signaling [58]. The presence of higher β2M in adult platelets would predict a greater ability of adult compared to neonatal platelets to induce a monocyte pro-inflammatory phenotype. Indeed, a recent murine study showed that transfusing pups with adult, but not neonatal, platelets triggered a pro-migratory monocyte phenotype that could potentially enhance tissue-level inflammation [56].

P-selectin, encoded by the *Selp* gene, is a component of the platelet alpha granules that translocates to the platelet surface upon platelet activation, making it available to bind its receptor (PSGL-1) located on monocytes and neutrophils. In this way, platelet P-selectin regulates monocyte functions, particularly pro-inflammatory IL-8 and MCP-1 cytokine production [59, 60]. *Selp* expression is developmentally regulated in mice, with neonatal megakaryocytes [61] and platelets [56] having significantly lower *Selp* mRNA levels compared to their adult counterparts. We found no significant differences in total P-selectin protein levels between resting neonatal and adult human platelets, consistent with a prior study that measured P-selectin in human adult and neonatal platelets by western blot [49]. This suggests species-related differences in the developmental regulation of this molecule. However, *activated* human neonatal platelets have significantly lower surface P-selectin exposure compared to *activated* adult platelets, due to intracellular signaling defects and decreased degranulation [49]. Thus, the phenotype of *activated* neonatal platelets in regard to P-selectin exposure is similar in both species, although the underlying mechanisms seem to be different.

Platelet derived growth factor (PDGF) is a member of the human growth factor family that is contained in the platelets alpha granules and released upon platelet activation. Several studies have shown that PDGF strongly contributes to airway remodeling in asthma, both by inducing the migration and proliferation of airway smooth muscle cells and by increasing the synthesis of collagen by fibroblasts [62]. Increased PDGF subunits in adult platelets could help explain the higher incidence of bronchopulmonary dysplasia (BPD) among preterm infants exposed to more platelet transfusions, since this is a disease process also characterized by airway obstruction and lung fibrosis [32, 63].

Neonatal and adult samples also clustered separately in our phosphoproteomics analysis, with 1183 phosphoproteins present in significantly different abundance. Pathway analysis of the 445 phosphoproteins that were significantly more abundant in adult platelets showed enrichment in enzyme regulation, particularly of GTPase enzymes. This is consistent with the known higher responsiveness of adult platelets to most agonists. GTPases act as signaling switches in signal transduction from platelet surface receptors to intracellular pathways, which mediate platelet activation [64]. Prior studies have shown developmental differences in platelet activation in response to the thromboxane analogue U46619 due to decreased signal transduction in neonatal platelets [65]. Of particular interest was the presence of phosphorylated RAP1GAP2 *exclusively* in adult platelets. Rap1GAP2 is a highly phosphorylated GTPase-activating protein that inhibits Rap1, a small guanine-nucleotide-binding protein that tightly regulates integrin activation in platelets [66]. In addition, Rap1GAP2 also binds to synaptotagmin-like protein 1 (Slp1) to regulate platelet dense granule secretion [67]. These findings suggest important links between altered RAP1GAP2 activities and adult platelet activation and/or degranulation.

Insulin growth factor signaling phosphopeptides from IGF1R, IGF2BP1 and IGF2BP3 were detected only in neonatal platelets. This likely reflects the importance of IGF-2 signaling in fetal and neonatal, but not adult, megakaryocytes [68, 69]. Insulin-like Growth Factor 2 mRNA-Binding Protein 3 (IGF2BP3) was also significantly more abundant at the total protein and phosphoprotein levels in neonatal compared to adult platelets (3.7-fold, p_adj_=3.3×10^-5^ for total protein, 5.6-fold, p_adj_=1.6×10^-4^ for S438-phosphopeptide, **Figure 5C, Table S2**, and **Table S6**). IGF2BP3 is known to regulate the fetal megakaryocyte program and is present at significantly higher concentrations in neonatal compared to adult megakaryocytes [55]. Evaluation of the role of enhanced IGF2BP3 phosphorylation in neonatal platelets warrants further study.

Our findings also revealed factors that could underlie differences in neonatal vs adult platelet degranulation [27]. For example, phosphorylated MARCKS was also more abundant in resting neonatal platelets and has been associated with calcium signaling regulation (**Figure 5C-D**). MARCKS and MARCKS-derived peptides bind to membrane lipids in megakaryocytes and platelets [70, 71] and have been implicated in platelet serotonin release [72]. MARCKS also regulates leukocyte degranulation [73], and MARCKS-derived peptides antagonize coagulation *in vivo* by inhibiting interactions between factor Xa and phosphatidylserine residues [71]. Future experiments will test whether increased MARCKS abundance and/or phosphorylation can mechanistically explain the recently described dissociation between degranulation and GPIIb/IIIa conformational change in activated neonatal platelets [74].

There were some limitations to our study design. First, we noted mild platelet activation during platelet isolation (**Figure 1**). This was an unavoidable consequence of platelet processing, which is important to recognize in order to provide context for potential changes in protein-based signaling or activation-related phosphorylation activities [75]. However, our samples clustered well based on developmental status irrespective of P-selectin MFI, and analyses performed after excluding potential outliers did not substantively change the results (data not shown). Second, umbilical cord blood and adult peripheral blood represent markedly different platelet sources. It was necessary to collect cord blood to obtain enough material, since we could not have purified an adequate number of platelets from infant peripheral blood to perform our study. Adult peripheral blood and umbilical cord blood also represent very different plasma environments for platelets, with altered immune-related activities or other variables that could affect platelet content or signaling. Indeed, we noted differences between adult and neonatal platelets in multiple plasma proteins, including complement factors (**Figure 3**). Given the presence of an open canalicular system in platelets, it is likely that some plasma proteins are retained inside platelets despite thorough washing. Thus, studying the platelet proteome may also provide a window into developmental differences in plasma proteins.

In conclusion, our study identified significant differences in protein content and phosphorylation status between neonatal vs adult platelets. Adult and neonatal platelets are markedly different cells with distinct functions that are tailored for their unique and developmentally regulated hemostatic and immune systems. The presence of increased complement and immunomodulatory proteins in adult platelets might help explain some of the deleterious effects associated with adult platelet transfusions in vulnerable neonates and infants.

## Acknowledgments

We thank Dr. Henry A. Feldman, PhD (Boston Children’s Hospital) for statistical advice and expertise. This project was funded by the National Institutes of Health (K99-HL156052 to CST, K99-HL156051 to PD, P01-HL046925 to MSV).

## Author Contributions

CST, PD, and MSV conceived the project. CST, PD, HF, ZJL, HZ, HD, JR, LAS, HI, and MSV collected, analyzed, and/or interpreted data. CST, PD, and MSV wrote the paper. All authors reviewed and approved the final version of the manuscript.

## References

[1] Maouia A, Rebetz J, Kapur R, Semple JW. The Immune Nature of Platelets Revisited. Transfus Med Rev. 2020; 34: 209–20. 10.1016/j.tmrv.2020.09.005.

[2] Koupenova M, Livada AC, Morrell CN. Platelet and Megakaryocyte Roles in Innate and Adaptive Immunity. Circ Res. 2022; 130: 288–308. 10.1161/CIRCRESAHA.121.319821.

[3] Weyrich AS. Platelets: more than a sack of glue. Hematology Am Soc Hematol Educ Program. 2014; 2014: 400–3. 2014/1/400 [pii]10.1182/asheducation-2014.1.400 [doi].

[4] Gay LJ, Felding-Habermann B. Contribution of platelets to tumour metastasis. Nat Rev Cancer. 2011; 11: 123–34. 10.1038/nrc3004.

[5] Sierko E, Wojtukiewicz MZ. Platelets and angiogenesis in malignancy. Semin Thromb Hemost. 2004; 30: 95-108. 10.1055/s-2004-822974.

[6] Battinelli EM, Markens BA, Italiano JE, Jr. Release of angiogenesis regulatory proteins from platelet alpha granules: modulation of physiologic and pathologic angiogenesis. Blood. 2011; 118: 1359–69. 10.1182/blood-2011-02-334524.

[7] Cognasse F, Nguyen KA, Damien P, McNicol A, Pozzetto B, Hamzeh-Cognasse H, Garraud O. The Inflammatory Role of Platelets via Their TLRs and Siglec Receptors. Front Immunol. 2015; 6: 83. 10.3389/fimmu.2015.00083.

[8] Suzuki-Inoue K, Fuller GL, Garcia A, Eble JA, Pohlmann S, Inoue O, Gartner TK, Hughan SC, Pearce AC, Laing GD, Theakston RD, Schweighoffer E, Zitzmann N, Morita T, Tybulewicz VL, Ozaki Y, Watson SP. A novel Syk-dependent mechanism of platelet activation by the C-type lectin receptor CLEC-2. Blood. 2006; 107: 542–9. 10.1182/blood-2005-05-1994.

[9] Zhang S, Zhang S, Hu L, Zhai L, Xue R, Ye J, Chen L, Cheng G, Mruk J, Kunapuli SP, Ding Z. Nucleotide-binding oligomerization domain 2 receptor is expressed in platelets and enhances platelet activation and thrombosis. Circulation. 2015; 131: 1160–70. 10.1161/CIRCULATIONAHA.114.013743.

[10] Elzey BD, Tian J, Jensen RJ, Swanson AK, Lees JR, Lentz SR, Stein CS, Nieswandt B, Wang Y, Davidson BL, Ratliff TL. Platelet-mediated modulation of adaptive immunity. A communication link between innate and adaptive immune compartments. Immunity. 2003; 19: 9–19. 10.1016/s1074-7613(03)00177-8.

[11] Kornerup KN, Salmon GP, Pitchford SC, Liu WL, Page CP. Circulating platelet-neutrophil complexes are important for subsequent neutrophil activation and migration. J Appl Physiol (1985). 2010; 109: 758–67. 10.1152/japplphysiol.01086.2009.

[12] Freedman JE, Loscalzo J. Platelet-monocyte aggregates: bridging thrombosis and inflammation. Circulation. 2002; 105: 2130–2. 10.1161/01.cir.0000017140.26466.f5.

[13] Lisman T. Platelet-neutrophil interactions as drivers of inflammatory and thrombotic disease. Cell Tissue Res. 2018; 371: 567–76. 10.1007/s00441-017-2727-4.

[14] Lam FW, Vijayan KV, Rumbaut RE. Platelets and Their Interactions with Other Immune Cells. Compr Physiol. 2015; 5: 1265–80. 10.1002/cphy.c140074.

[15] Clark SR, Ma AC, Tavener SA, McDonald B, Goodarzi Z, Kelly MM, Patel KD, Chakrabarti S, McAvoy E, Sinclair GD, Keys EM, Allen-Vercoe E, Devinney R, Doig CJ, Green FH, Kubes P. Platelet TLR4 activates neutrophil extracellular traps to ensnare bacteria in septic blood. Nat Med. 2007; 13: 463–9. nm1565 [pii]10.1038/nm1565.

[16] Del Conde I, Cruz MA, Zhang H, Lopez JA, Afshar-Kharghan V. Platelet activation leads to activation and propagation of the complement system. J Exp Med. 2005; 201: 871–9. 10.1084/jem.20041497.

[17] Peerschke EI, Yin W, Grigg SE, Ghebrehiwet B. Blood platelets activate the classical pathway of human complement. J Thromb Haemost. 2006; 4: 2035–42. 10.1111/j.1538-7836.2006.02065.x.

[18] Wiedmer T, Esmon CT, Sims PJ. Complement proteins C5b-9 stimulate procoagulant activity through platelet prothrombinase. Blood. 1986; 68: 875–80.

[19] Gaertner F, Ahmad Z, Rosenberger G, Fan S, Nicolai L, Busch B, Yavuz G, Luckner M, Ishikawa-Ankerhold H, Hennel R, Benechet A, Lorenz M, Chandraratne S, Schubert I, Helmer S, Striednig B, Stark K, Janko M, Bottcher RT, Verschoor A, Leon C, Gachet C, Gudermann T, Mederos YSM, Pincus Z, Iannacone M, Haas R, Wanner G, Lauber K, Sixt M, Massberg S. Migrating Platelets Are Mechano-scavengers that Collect and Bundle Bacteria. Cell. 2017; 171: 1368–82 e23. 10.1016/j.cell.2017.11.001.

[20] Palankar R, Kohler TP, Krauel K, Wesche J, Hammerschmidt S, Greinacher A. Platelets kill bacteria by bridging innate and adaptive immunity via platelet factor 4 and FcgammaRIIA. J Thromb Haemost. 2018; 16: 1187–97. 10.1111/jth.13955.

[21] Golebiewska EM, Poole AW. Platelet secretion: From haemostasis to wound healing and beyond. Blood Rev. 2015; 29: 153–62. 10.1016/j.blre.2014.10.003.

[22] Kao KJ, Cook DJ, Scornik JC. Quantitative analysis of platelet surface HLA by W6/32 anti-HLA monoclonal antibody. Blood. 1986; 68: 627–32.

[23] Denis MM, Tolley ND, Bunting M, Schwertz H, Jiang H, Lindemann S, Yost CC, Rubner FJ, Albertine KH, Swoboda KJ, Fratto CM, Tolley E, Kraiss LW, McIntyre TM, Zimmerman GA, Weyrich AS. Escaping the nuclear confines: signal-dependent pre-mRNA splicing in anucleate platelets. Cell. 2005; 122: 379–91. 10.1016/j.cell.2005.06.015.

[24] Kieffer N, Guichard J, Farcet JP, Vainchenker W, Breton-Gorius J. Biosynthesis of major platelet proteins in human blood platelets. Eur J Biochem. 1987; 164: 189–95. 10.1111/j.1432-1033.1987.tb11010.x.

[25] Lindemann S, Tolley ND, Dixon DA, McIntyre TM, Prescott SM, Zimmerman GA, Weyrich AS. Activated platelets mediate inflammatory signaling by regulated interleukin 1beta synthesis. J Cell Biol. 2001; 154: 485–90. 10.1083/jcb.200105058 [doi]154/3/485 [pii].

[26] Weyrich AS, Schwertz H, Kraiss LW, Zimmerman GA. Protein synthesis by platelets: historical and new perspectives. J Thromb Haemost. 2009; 7: 241–6. 10.1111/j.1538-7836.2008.03211.x.

[27] Ferrer-Marin F, Sola-Visner M. Neonatal platelet physiology and implications for transfusion. Platelets. 2021: 1–9. 10.1080/09537104.2021.1962837.

[28] Ferrer-Marin F, Stanworth S, Josephson C, Sola-Visner M. Distinct differences in platelet production and function between neonates and adults: implications for platelet transfusion practice. Transfusion. 2013; 53: 2814–21; quiz 3. 10.1111/trf.12343.

[29] Toulon P. Developmental hemostasis: laboratory and clinical implications. Int J Lab Hematol. 2016; 38 Suppl 1: 66–77. 10.1111/ijlh.12531.

[30] Roschitz B, Sudi K, Kostenberger M, Muntean W. Shorter PFA-100 closure times in neonates than in adults: role of red cells, white cells, platelets and von Willebrand factor. Acta Paediatr. 2001; 90: 664–70.

[31] Saxonhouse MA, Garner R, Mammel L, Li Q, Muller KE, Greywoode J, Miller C, Sola-Visner M. Closure times measured by the platelet function analyzer PFA-100 are longer in neonatal blood compared to cord blood samples. Neonatology. 2010; 97: 242–9. 10.1159/000253755.

[32] Curley A, Stanworth SJ, Willoughby K, Fustolo-Gunnink SF, Venkatesh V, Hudson C, Deary A, Hodge R, Hopkins V, Lopez Santamaria B, Mora A, Llewelyn C, D’Amore A, Khan R, Onland W, Lopriore E, Fijnvandraat K, New H, Clarke P, Watts T, PlaNe TMC. Randomized Trial of Platelet-Transfusion Thresholds in Neonates. N Engl J Med. 2019; 380: 242–51. 10.1056/NEJMoa1807320.

[33] Caparros-Perez E, Teruel-Montoya R, Lopez-Andreo MJ, Llanos MC, Rivera J, Palma-Barqueros V, Blanco JE, Vicente V, Martinez C, Ferrer-Marin F. Comprehensive comparison of neonate and adult human platelet transcriptomes. PLoS One. 2017; 12: e0183042. 10.1371/journal.pone.0183042.

[34] Liu Z, Avila C, Malone LE, Gnatenko DV, Sheriff J, Zhu W, Bahou WF. Age-restricted functional and developmental differences of neonatal platelets. J Thromb Haemost. 2022; 20: 2632–45. 10.1111/jth.15847.

[35] Stokhuijzen E, Koornneef JM, Nota B, van den Eshof BL, van Alphen FPJ, van den Biggelaar M, van der Zwaan C, Kuijk C, Mertens K, Fijnvandraat K, Meijer AB. Differences between Platelets Derived from Neonatal Cord Blood and Adult Peripheral Blood Assessed by Mass Spectrometry. J Proteome Res. 2017; 16: 3567–75. 10.1021/acs.jproteome.7b00298.

[36] Mertins P, Qiao JW, Patel J, Udeshi ND, Clauser KR, Mani DR, Burgess MW, Gillette MA, Jaffe JD, Carr SA. Integrated proteomic analysis of post-translational modifications by serial enrichment. Nat Methods. 2013; 10: 634–7. 10.1038/nmeth.2518.

[37] Dominguez S, Rodriguez G, Fazelinia H, Ding H, Spruce L, Seeholzer SH, Dong H. Sex Differences in the Phosphoproteomic Profiles of APP/PS1 Mice after Chronic Unpredictable Mild Stress. J Alzheimers Dis. 2020; 74: 1131–42. 10.3233/JAD-191009.

[38] McNulty DE, Annan RS. Hydrophilic interaction chromatography reduces the complexity of the phosphoproteome and improves global phosphopeptide isolation and detection. Mol Cell Proteomics. 2008; 7: 971–80. 10.1074/mcp.M700543-MCP200.

[39] Tyanova S, Temu T, Cox J. The MaxQuant computational platform for mass spectrometry-based shotgun proteomics. Nat Protoc. 2016; 11: 2301–19. 10.1038/nprot.2016.136.

[40] Bruderer R, Bernhardt OM, Gandhi T, Miladinovic SM, Cheng LY, Messner S, Ehrenberger T, Zanotelli V, Butscheid Y, Escher C, Vitek O, Rinner O, Reiter L. Extending the limits of quantitative proteome profiling with data-independent acquisition and application to acetaminophen-treated three-dimensional liver microtissues. Mol Cell Proteomics. 2015; 14: 1400–10. 10.1074/mcp.M114.044305.

[41] Mi H, Muruganujan A, Ebert D, Huang X, Thomas PD. PANTHER version 14: more genomes, a new PANTHER GO-slim and improvements in enrichment analysis tools. Nucleic Acids Res. 2019; 47: D419–D26. 10.1093/nar/gky1038.

[42] Legeay M, Doncheva NT, Morris JH, Jensen LJ. Visualize omics data on networks with Omics Visualizer, a Cytoscape App. F1000Res. 2020; 9: 157. 10.12688/f1000research.22280.2.

[43] Burkhart JM, Vaudel M, Gambaryan S, Radau S, Walter U, Martens L, Geiger J, Sickmann A, Zahedi RP. The first comprehensive and quantitative analysis of human platelet protein composition allows the comparative analysis of structural and functional pathways. Blood. 2012; 120: e73–82. 10.1182/blood-2012-04-416594.

[44] Solari FA, Krahn D, Swieringa F, Verhelst S, Rassaf T, Tasdogan A, Zahedi RP, Lorenz K, Renne T, Heemskerk JWM, Sickmann A. Multi-omics approaches to study platelet mechanisms. Curr Opin Chem Biol. 2023; 73: 102253. 10.1016/j.cbpa.2022.102253.

[45] Mann M, Ong SE, Gronborg M, Steen H, Jensen ON, Pandey A. Analysis of protein phosphorylation using mass spectrometry: deciphering the phosphoproteome. Trends Biotechnol. 2002; 20: 261–8. 10.1016/s0167-7799(02)01944-3.

[46] Gunawardena J. Multisite protein phosphorylation makes a good threshold but can be a poor switch. Proc Natl Acad Sci U S A. 2005; 102: 14617–22. 10.1073/pnas.0507322102.

[47] Stalker TJ, Newman DK, Ma P, Wannemacher KM, Brass LF. Platelet signaling. Handb Exp Pharmacol. 2012: 59-85. 10.1007/978-3-642-29423-5_3.

[48] Yang YS, Strittmatter SM. The reticulons: a family of proteins with diverse functions. Genome Biol. 2007; 8: 234. 10.1186/gb-2007-8-12-234.

[49] Caparros-Perez E, Teruel-Montoya R, Palma-Barquero V, Torregrosa JM, Blanco JE, Delgado JL, Lozano ML, Vicente V, Sola-Visner M, Rivera J, Martinez C, Ferrer-Marin F. Down Regulation of the Munc18b-syntaxin-11 Complex and beta1-tubulin Impairs Secretion and Spreading in Neonatal Platelets. Thromb Haemost. 2017; 117: 2079–91. 10.1160/TH17-04-0241.

[50] Kuleshov MV, Xie Z, London ABK, Yang J, Evangelista JE, Lachmann A, Shu I, Torre D, Ma’ayan A. KEA3: improved kinase enrichment analysis via data integration. Nucleic Acids Res. 2021; 49: W304–W16. 10.1093/nar/gkab359.

[51] Senis YA, Mazharian A, Mori J. Src family kinases: at the forefront of platelet activation. Blood. 2014; 124: 2013–24. 10.1182/blood-2014-01-453134.

[52] Liu ZJ, Italiano J, Jr., Ferrer-Marin F, Gutti R, Bailey M, Poterjoy B, Rimsza L, Sola-Visner M. Developmental differences in megakaryocytopoiesis are associated with up-regulated TPO signaling through mTOR and elevated GATA-1 levels in neonatal megakaryocytes. Blood. 2011; 117: 4106–17. blood-2010-07-293092 [pii] 10.1182/blood-2010-07-293092.

[53] Guo L, Shen S, Rowley JW, Tolley ND, Jia W, Manne BK, McComas KN, Bolingbroke B, Kosaka Y, Krauel K, Denorme F, Jacob SP, Eustes AS, Campbell RA, Middleton EA, He X, Brown SM, Morrell CN, Weyrich AS, Rondina MT. Platelet MHC class I mediates CD8+ T-cell suppression during sepsis. Blood. 2021; 138: 401–16. 10.1182/blood.2020008958.

[54] Koupenova M, Clancy L, Corkrey HA, Freedman JE. Circulating Platelets as Mediators of Immunity, Inflammation, and Thrombosis. Circ Res. 2018; 122: 337–51. 10.1161/CIRCRESAHA.117.310795.

[55] Elagib KE, Lu CH, Mosoyan G, Khalil S, Zasadzinska E, Foltz DR, Balogh P, Gru AA, Fuchs DA, Rimsza LM, Verhoeyen E, Sanso M, Fisher RP, Iancu-Rubin C, Goldfarb AN. Neonatal expression of RNA-binding protein IGF2BP3 regulates the human fetal-adult megakaryocyte transition. J Clin Invest. 2017; 127: 2365–77. 10.1172/JCI88936.

[56] Maurya P, Ture SK, Li C, Scheible KM, McGrath KE, Palis J, Morrell CN. Transfusion of Adult, but Not Neonatal, Platelets Promotes Monocyte Trafficking in Neonatal Mice. Arterioscler Thromb Vasc Biol. 2023. 10.1161/ATVBAHA.122.318162.

[57] Woo AJ, Wieland K, Huang H, Akie TE, Piers T, Kim J, Cantor AB. Developmental differences in IFN signaling affect GATA1s-induced megakaryocyte hyperproliferation. J Clin Invest. 2013. 40609 [pii]10.1172/JCI40609.

[58] Hilt ZT, Pariser DN, Ture SK, Mohan A, Quijada P, Asante AA, Cameron SJ, Sterling JA, Merkel AR, Johanson AL, Jenkins JL, Small EM, McGrath KE, Palis J, Elliott MR, Morrell CN. Platelet-derived beta2M regulates monocyte inflammatory responses. JCI Insight. 2019; 4. 10.1172/jci.insight.122943.

[59] Weyrich AS, Elstad MR, McEver RP, McIntyre TM, Moore KL, Morrissey JH, Prescott SM, Zimmerman GA. Activated platelets signal chemokine synthesis by human monocytes. J Clin Invest. 1996; 97: 1525–34. 10.1172/JCI118575.

[60] Weyrich AS, McIntyre TM, McEver RP, Prescott SM, Zimmerman GA. Monocyte tethering by P-selectin regulates monocyte chemotactic protein-1 and tumor necrosis factor-alpha secretion. Signal integration and NF-kappa B translocation. J Clin Invest. 1995; 95: 2297–303. 10.1172/JCI117921.

[61] Palis J SM, McGrath KE, Kingsley PD. P-selectin expression and platelet function are developmentally regulated. Exp Hematol. 2015; 43: S87.

[62] Kardas G, Daszynska-Kardas A, Marynowski M, Brzakalska O, Kuna P, Panek M. Role of Platelet-Derived Growth Factor (PDGF) in Asthma as an Immunoregulatory Factor Mediating Airway Remodeling and Possible Pharmacological Target. Front Pharmacol. 2020; 11: 47. 10.3389/fphar.2020.00047.

[63] Moore CM, D’Amore A, Fustolo-Gunnink S, Hudson C, Newton A, Santamaria BL, Deary A, Hodge R, Hopkins V, Mora A, Llewelyn C, Venkatesh V, Khan R, Willoughby K, Onland W, Fijnvandraat K, New HV, Clarke P, Lopriore E, Watts T, Stanworth S, Curley A, PlaNe TM. Two-year outcomes following a randomised platelet transfusion trial in preterm infants. Arch Dis Child Fetal Neonatal Ed. 2023. 10.1136/archdischild-2022-324915.

[64] Comer SP. Turning Platelets Off and On: Role of RhoGAPs and RhoGEFs in Platelet Activity. Front Cardiovasc Med. 2021; 8: 820945. 10.3389/fcvm.2021.820945.

[65] Israels SJ, Odaibo FS, Robertson C, McMillan EM, McNicol A. Deficient thromboxane synthesis and response in platelets from premature infants. Pediatr Res. 1997; 41: 218–23.

[66] Schultess J, Danielewski O, Smolenski AP. Rap1GAP2 is a new GTPase-activating protein of Rap1 expressed in human platelets. Blood. 2005; 105: 3185–92. 10.1182/blood-2004-09-3605.

[67] Neumuller O, Hoffmeister M, Babica J, Prelle C, Gegenbauer K, Smolenski AP. Synaptotagmin-like protein 1 interacts with the GTPase-activating protein Rap1GAP2 and regulates dense granule secretion in platelets. Blood. 2009; 114: 1396–404. 10.1182/blood-2008-05-155234.

[68] Zhang CC, Lodish HF. Insulin-like growth factor 2 expressed in a novel fetal liver cell population is a growth factor for hematopoietic stem cells. Blood. 2004; 103: 2513–21. 10.1182/blood-2003-08-2955.

[69] Klusmann JH, Godinho FJ, Heitmann K, Maroz A, Koch ML, Reinhardt D, Orkin SH, Li Z. Developmental stage-specific interplay of GATA1 and IGF signaling in fetal megakaryopoiesis and leukemogenesis. Genes Dev. 2010; 24: 1659–72. 24/15/1659 [pii]10.1101/gad.1903410.

[70] Machlus KR, Wu SK, Stumpo DJ, Soussou TS, Paul DS, Campbell RA, Kalwa H, Michel T, Bergmeier W, Weyrich AS, Blackshear PJ, Hartwig JH, Italiano JE, Jr. Synthesis and dephosphorylation of MARCKS in the late stages of megakaryocyte maturation drive proplatelet formation. Blood. 2016; 127: 1468–80. 10.1182/blood-2015-08-663146.

[71] Kastelowitz N, Tamura R, Onasoga A, Stalker TJ, White OR, Brown PN, Brodsky GL, Brass LF, Branchford BR, Di Paola J, Yin H. Peptides derived from MARCKS block coagulation complex assembly on phosphatidylserine. Sci Rep. 2017; 7: 4275. 10.1038/s41598-017-04494-y.

[72] Elzagallaai A, Rose SD, Trifaro JM. Platelet secretion induced by phorbol esters stimulation is mediated through phosphorylation of MARCKS: a MARCKS-derived peptide blocks MARCKS phosphorylation and serotonin release without affecting pleckstrin phosphorylation. Blood. 2000; 95: 894–902.

[73] Takashi S, Park J, Fang S, Koyama S, Parikh I, Adler KB. A peptide against the N-terminus of myristoylated alanine-rich C kinase substrate inhibits degranulation of human leukocytes in vitro. Am J Respir Cell Mol Biol. 2006; 34: 647–52. 10.1165/rcmb.2006-0030RC.

[74] Weiss LJ, Drayss M, Mott K, Beck S, Unsin D, Just B, Speer CP, Hartel C, Andres O, Schulze H. Ontogenesis of functional platelet subpopulations from preterm and term neonates to adulthood: The PLINIUS study. Blood Adv. 2023. 10.1182/bloodadvances.2023009824.

[75] Huang J, Zhang P, Solari FA, Sickmann A, Garcia A, Jurk K, Heemskerk JWM. Molecular Proteomics and Signalling of Human Platelets in Health and Disease. Int J Mol Sci. 2021; 22. 10.3390/ijms22189860.

