## Supplementary Information for "Phosphoproteomics reveals content and signaling differences between neonatal and adult platelets"

**Supplemental Figures**

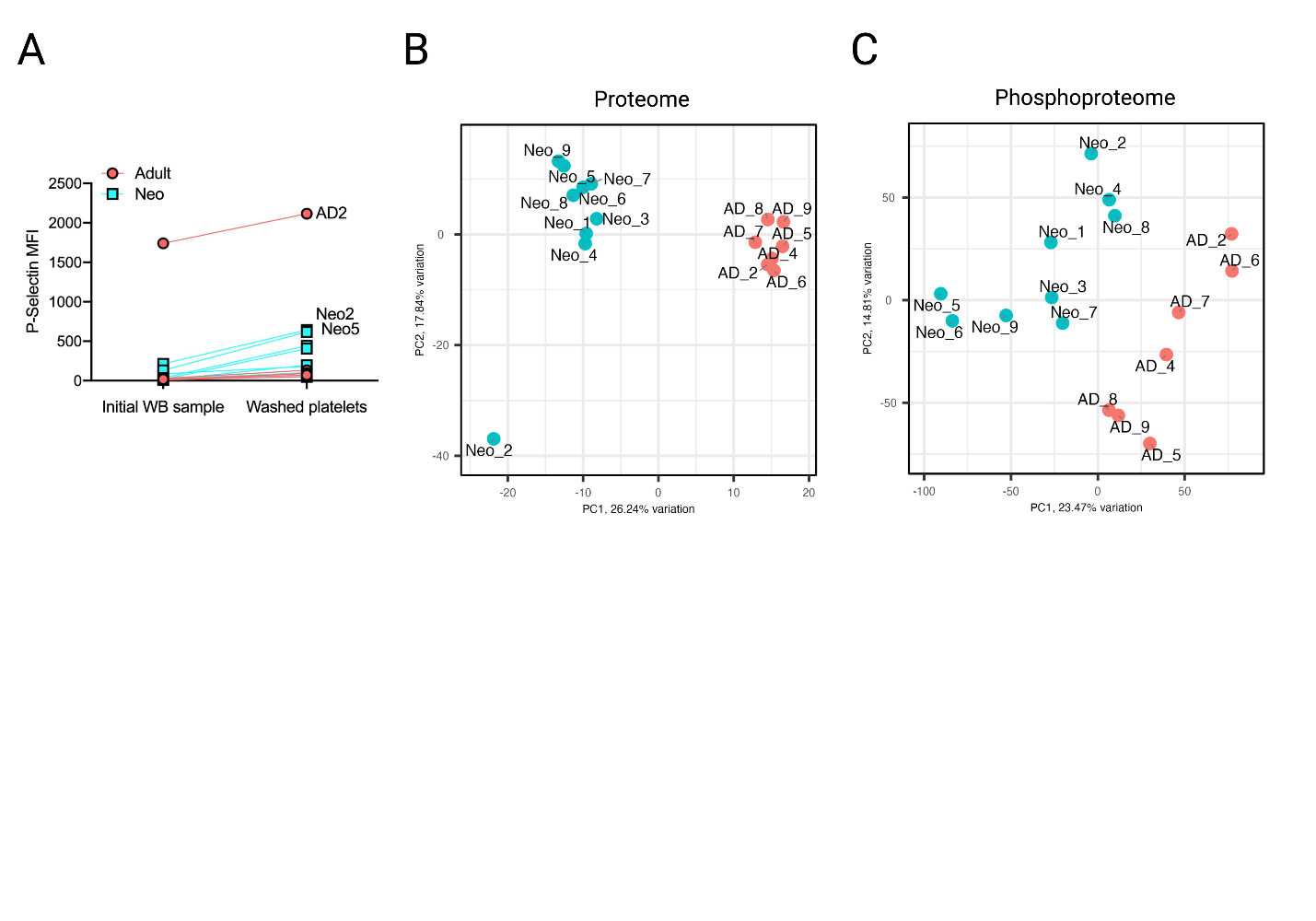

**Figure S1. Sample comparisons by P-selectin mean fluorescence intensity (MFI) and principal component analysis (PCA) plots. A.** P-selectin MFI of platelets in whole blood immediately after collection and following isolation and washing. Samples with highest P-selectin MFI are labeled. **B.** PCA plot for proteome analysis of neonatal and adult platelet samples with individual samples labeled. **C.** PCA plot for phosphoproteomic analysis with neonatal and adult platelet samples labeled. Note that sample AD2, an outlier by P-selectin MFI, clustered closely with other adult samples in PCA plots. Sample Neo2 was somewhat separate from other neonatal samples in the Proteome PCA plot, but sample Neo5, which had very similar P-selectin MFI after platelet isolation, clustered closely with the other neonatal samples.

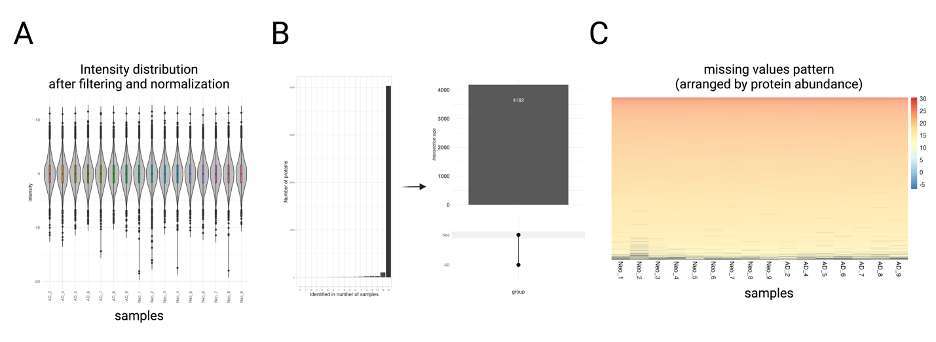

**Figure S2. Quality control and validation of neonatal and adult platelet proteome samples. A.** Intensity distribution plot after filtering and normalization of the data, with adult or neonatal samples shown across the bottom. **B.** Bar and upset plot showing that the majority of all high-confidence proteins identified in our study were present in all samples of both neonatal and adult origin, including 4182 high-confidence proteins that we took forward for further analysis. **C.** Missingness plot shows that most proteins identified in our analysis were found in all samples. Each row represents individual proteins, arranged by abundance and colored by the colored legend at right. Samples are shown across the bottom. Samples where individual proteins are missing are shown in black. These occur only in the least abundant proteins at the bottom of the plot.

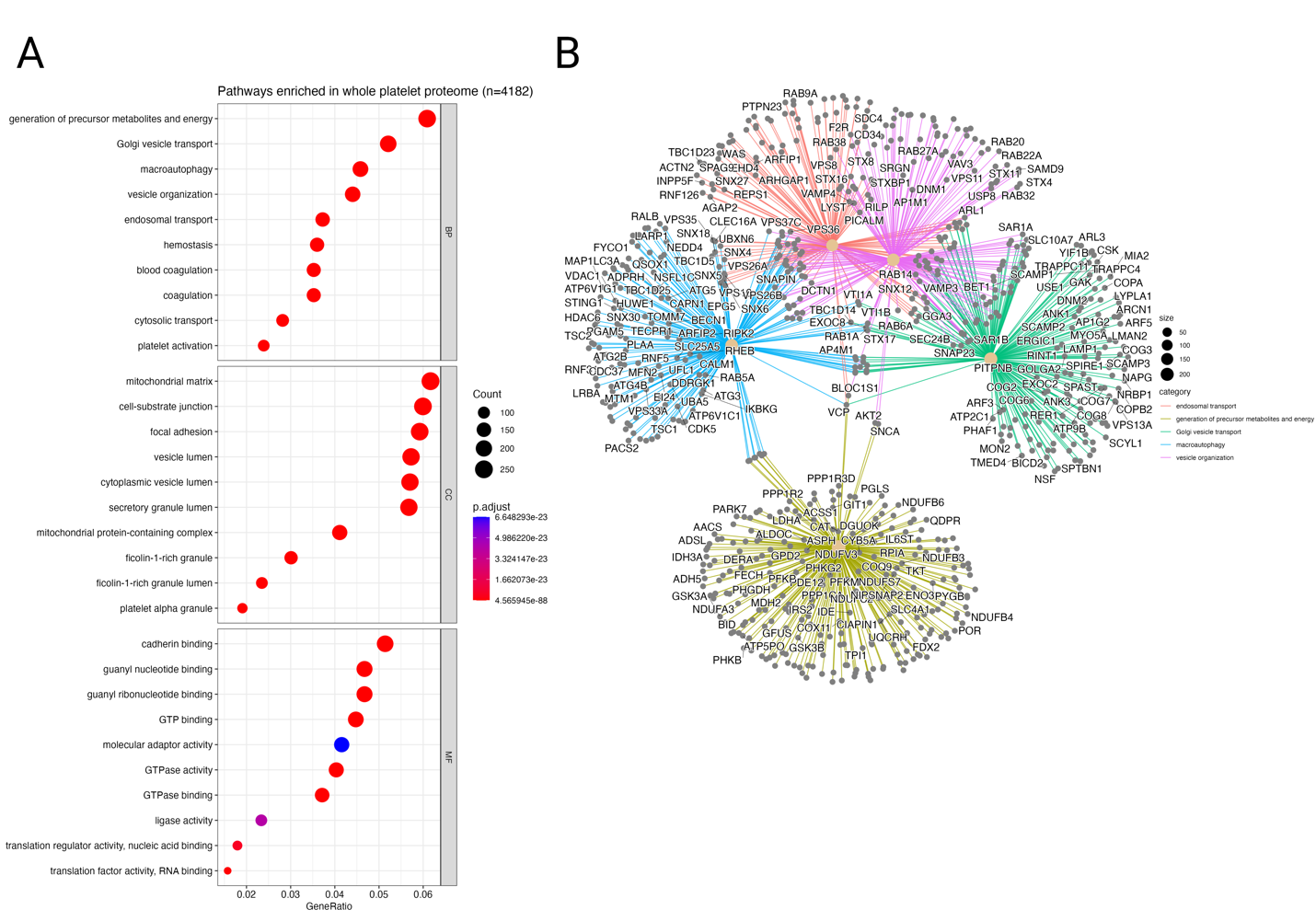

**Figure S3. Gene ontology enrichment for the platelet proteome, irrespective or neonatal or adult origin. A**. Dot plot showing pathway enrichment for the platelet proteome identified in our study (n=4182 proteins), as compared to the entire potential cell proteome/transcriptome. **B.** Web plot showing individual genes and interconnectedness of select pathways enriched in the platelet proteome. Pathway names are shown under ‘category’ and color coded.

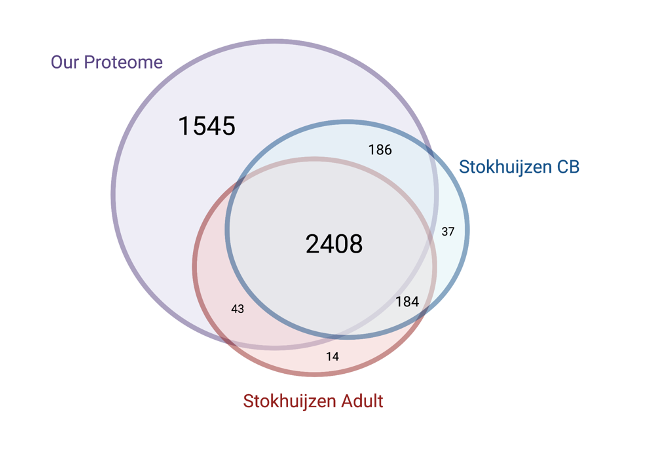

**Figure S4. Comparison of proteome results for our study and one prior study interrogating the neonatal vs adult platelet proteome.** Note that our neonatal/adult platelet proteome is encompassed by one circle since all 4182 proteins identified in our study were present in both neonatal and adult platelets. Stokhuijzen *et al* data were processed using the same pipeline as our data to facilitate comparison (**Methods**). This resulted in the identification of 2872 unique proteins.

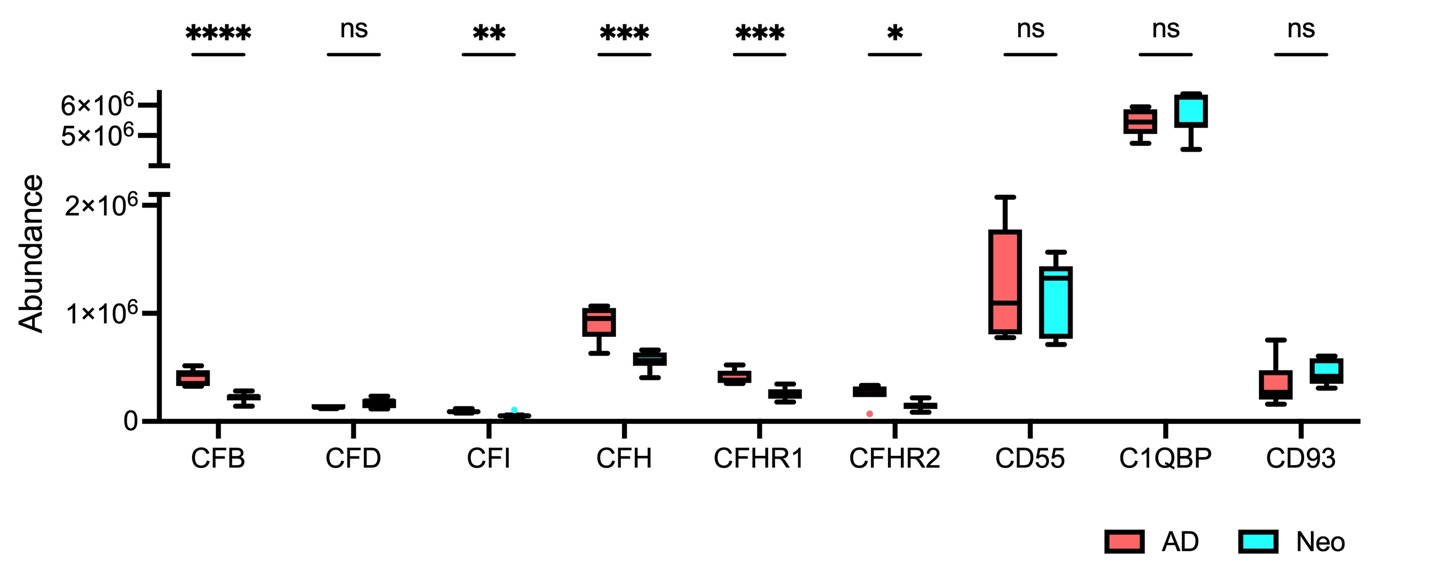

**Figure S5. Comparisons of total protein abundance for the indicated complement-related proteins.** Tukey box plots depict median with 25^th^-75^th^ interquartile range (IQR), and whiskers represent 1.5 times IQR. Statistical significance was d calculated using the Holm step-down procedure. ns, not significant. CFB, Complement factor B. CFD, Complement factor D. CFI, Complement factor I. CFH, Complement factor H. CFHR1, Complement factor H-related protein 1. CFHR2, Complement factor H-related protein 2. CD55, Complement decay-accelerating factor. C1QBP, Complement component 1 Q subcomponent-binding protein. CD93, Complement component C1Q receptor. *p<0.05, **p<0.01, ***p<0.001, ****p<0.0001.

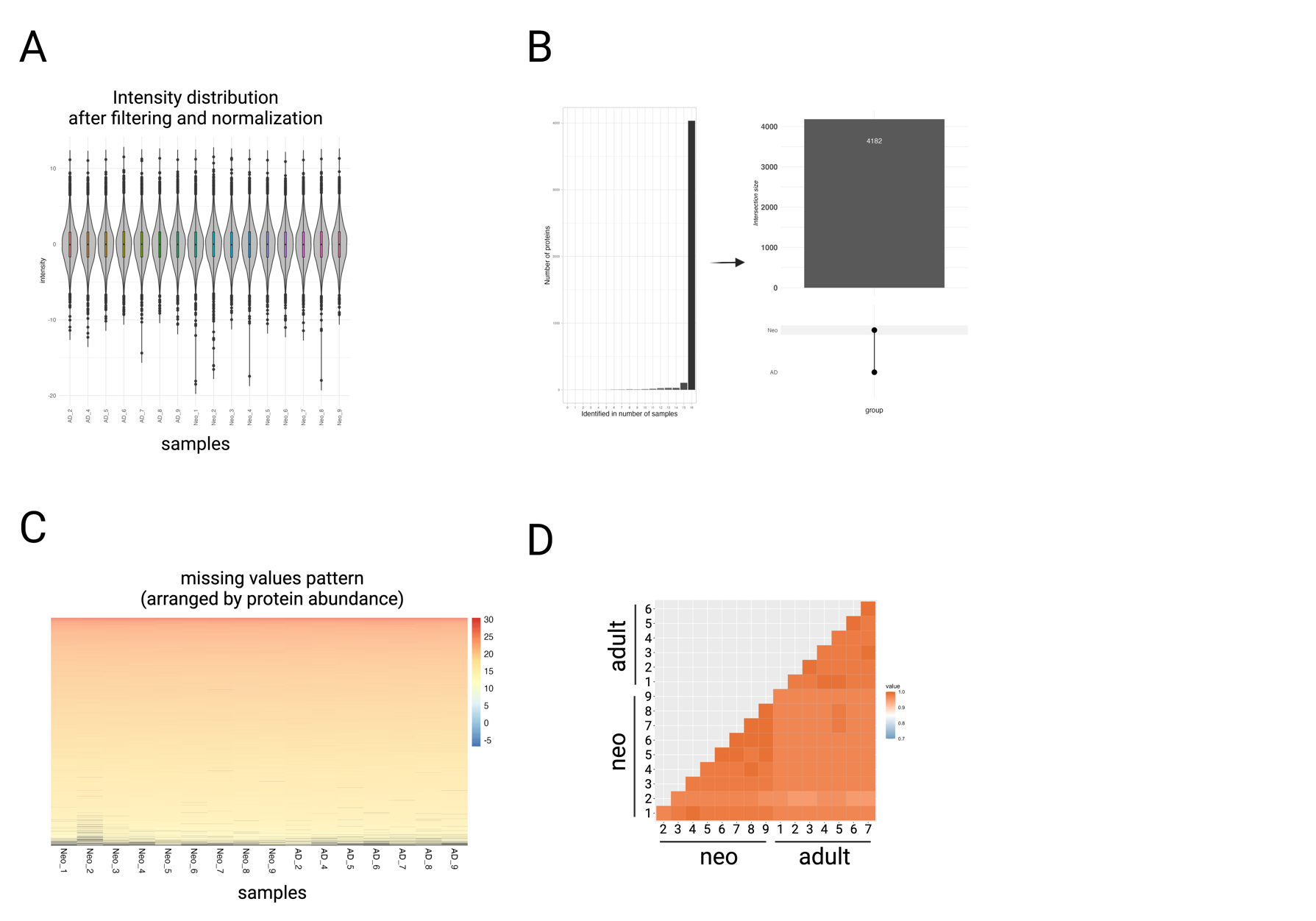

**Figure S6. Quality control and validation of neonatal and adult platelet phosphoproteome samples. A.** Intensity distribution plot after filtering and normalization of the data, with adult or neonatal samples shown across the bottom. **B.** Bar and upset plot showing that the majority of all high-confidence phosphopeptides identified in our study were present in all samples of both neonatal and adult origin, including 17,852 high-confidence phosphopeptides that we took forward for further analysis. Of these, 17,842 were found in both neonatal and adult samples whereas 8 were exclusive to neonatal platelets and 2 were exclusive to adult platelet samples. **C.** Missingness plot shows that most phosphopeptides identified in our analysis were found in all samples. Each row represents individual proteins, arranged by abundance and colored by the colored legend at right. Samples are shown across the bottom. Samples where individual proteins are missing are shown in black. These occurred in the least abundant phosphoproteins toward the bottom of the plot. **D**. Similarity plot comparing neonatal and adult platelet samples.

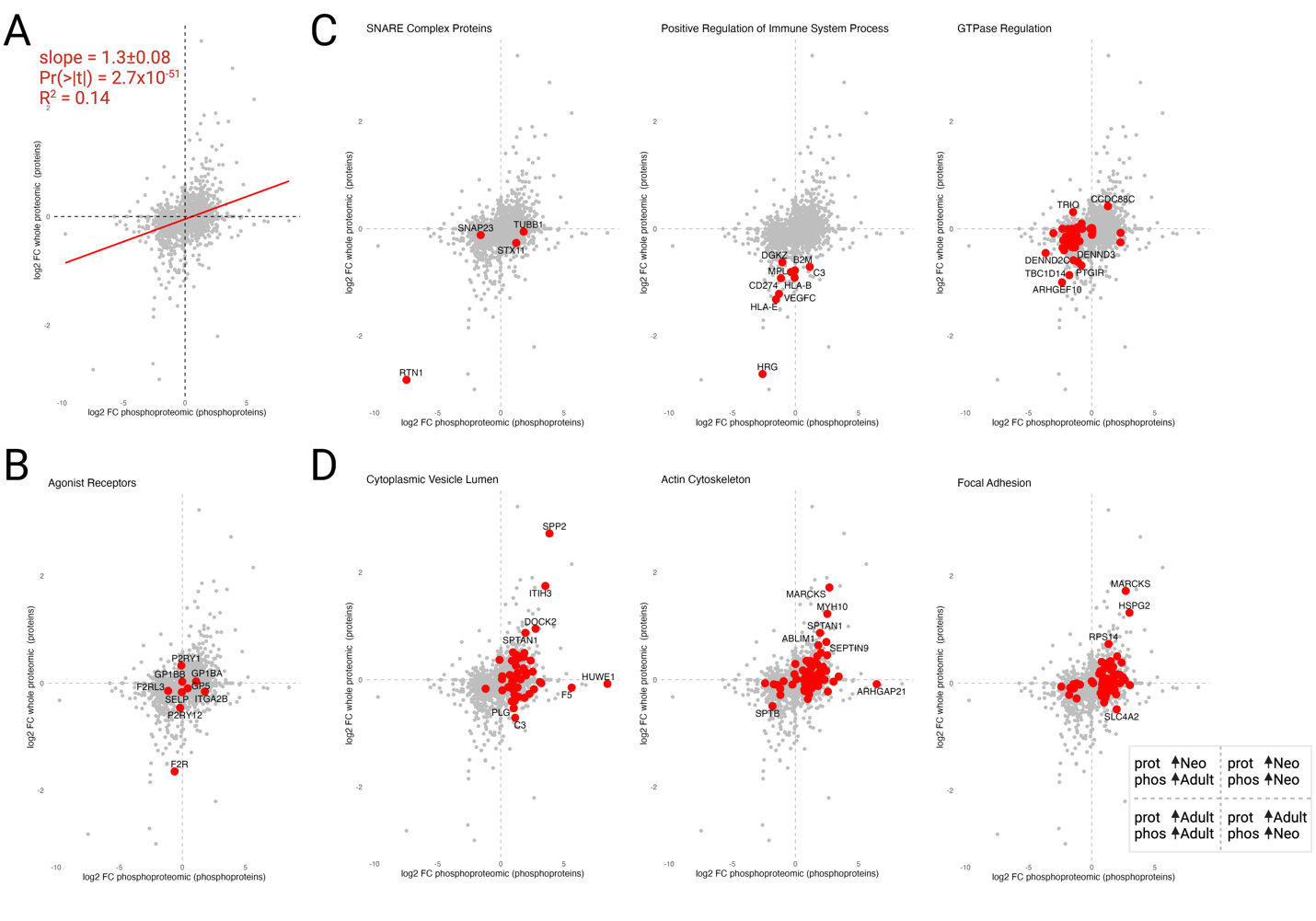

**Figure S7. Trends and pathway enrichment within our platelet proteome and phosphoproteome data. A.** Scatterplot depicting line of best fit and related statistics showing a positive correlation between protein abundance (y-axis) and phosphoprotein abundance (x-axis). **B.** Protein and phosphoprotein abundance for select agonist receptor proteins. **C.** Select pathways enriched in adult platelets based on our data and/or prior studies. **D.** Select pathways enriched in neonatal platelets based on our data.

**Supplemental Table Legends**

**Table S1.** R environment and packages used in this study.

**Table S2.** Protein enrichment in our proteomic analysis of resting platelets. The logFC [ log_2_(Fold Change) ] refers to the change in neonatal vs adult platelets, so a positive logFC reflects enrichment in neonatal platelets. Normalized average expression (AveExpr) is normalized to all detected proteins. Presented statistics include the t-statistic, p value (P.Value, unadjusted), adjusted p value (adj.P.Val, adjusted for multiple comparisons), and B-statistic (a measure of the log-odds ratio for differential expression).

**Table S3.** Gene ontology pathway enrichment for the platelet proteome compared to the entire human proteome/transcriptome. Column headers reflect related Ontology (BP=Biological Process, CC=Cell Compartment, MF=Molecular Function), Gene Ontology ID and Description, ratio of pathway-related proteins to the total protein number in our study (GeneRatio), expected ratio based on proteins in each pathway data set (BgRatio), p value (unadjusted), p value adjusted for multiple comparisons (p.adjust), q value (false discovery rate), identifications for related platelet proteins (geneID), and the total count of platelet proteins related to each data set. A total of 3974 out of 4182 proteins were recognized by the Gene Ontology database for these analyses.

**Table S4.** Gene ontology pathway enrichment for proteins enriched in adult vs neonatal platelets.

**Table S5.** Gene ontology pathway enrichment for proteins enriched in neonatal vs adult platelets.

**Table S6.** Phosphopeptide enrichment in our phosphoproteomic analysis of resting platelets. Rows represent data for individual phosphopeptides. In each row, the Protein and Uniprot identification are given alongside the specific peptide. The logFC [ log_2_(Fold Change) ] refers to the change in neonatal vs adult platelets, so a positive logFC reflects enrichment in neonatal platelets. Normalized average expression (AveExpr) is normalized to all detected proteins. Presented statistics include the t-statistic, p value (P.Value, unadjusted), adjusted p value (adj.P.Val, adjusted for multiple comparisons), and B-statistic (a measure of the log-odds ratio for differential expression).

**Table S7.** Gene ontology pathway enrichment for the entire platelet phosphoproteome (combined adult and neonatal data) compared to the platelet proteome defined in this study.

**Table S8.** Gene ontology pathway enrichment for phosphoproteins significantly more abundant in adult vs neonatal platelets compared to the platelet proteome defined in this study.

**Table S9.** Gene ontology pathway enrichment for phosphoproteins significantly more abundant in neonatal vs adult platelets compared to the platelet proteome defined in this study.
