## Supplementary Methods for "Phosphoproteomics reveals content and signaling differences between neonatal and adult platelets"

***Protein extraction.*** Platelet pellets were lysed in 8M urea lysis buffer (8 M urea, 75 mM NaCl, 50 mM Tris HCl pH 8.0), proteinase inhibitor cocktail (Roche, 11836170001) and PhosStop (Roche, 04906837001). Lysates were transferred to 2mL Eppendorf tubes containing 2 x 5mm stainless steel balls and placed into a ball mill (Retch, MM400) at 25 Hz and 30” per cycle for 6 cycles. The lysates were transferred to new tubes and centrifuged at 20,000g for 8 minutes at 4^o^C. Supernatants were then decanted and stored at -80^o^C [36]. The protein concentration of each supernatant was assessed by Micro BCA assay (Thermo Scientific).

***Protein hydrolysis.*** One milligram (1mg) protein from each sample was spiked with 1μg chicken ovalbumin (1:1000), which served as a phosphoprotein internal standard. Proteins were reduced with 5mM dithiothreitol for 45 min, followed by alkylation with 20 mM iodoacetamide for 45 min in the dark. Lysates were diluted with 50 mM Tris-HCl (pH 8.0) to reduce the total urea concentration to 0.8 M. To achieve proteolysis, samples were incubated with LysC (Wako,129-02541) at a 1:100 (enzyme:protein) ratio at 27ºC for 2 hrs, followed by trypsin (Promega, V5111) at a 1:50 (enzyme:protein) ratio at 27ºC overnight. Digests were acidified with formic acid (Thermo Scientific, 28905, final concentration 1%) and centrifuged at 20,000g for 10 min. Supernatants were desalted under a vacuum manifold using an Oasis HLB 96 well plate (60mg, Waters, 186000679) that had been preconditioned with 1 × 400μL 100% acetonitrile and equilibrated with 2 × 400μL 0.1% trifluoroacetic acid (J T Baker, 9470-00). Tryptic peptides were loaded into the plate, washed with 3 x 400μL 0.1% trifluoroacetic acid, and eluted into a protein Lobind plate 96/2000 uL (Eppendorf, EP0030504305) with 3 x 300 μL 50% acetonitrile/0.1% trifluoroacetic acid.

***Allocation of digests for further processing.*** Five percent (5%) of each sample was used for whole proteome analysis. 75% of each sample was used for IMAC phosphopeptide enrichment [36-38]. The remaining 20% for each sample was pooled and used for phosphopeptide spectral library generation. All peptides were lyophilized and stored at -80ºC. Peptides for spectral library generation were solubilized in 2% acetonitrile, 5 mM ammonium formate (pH 10, Sigma 17843) and separated by high pH reverse phase (RP)-HPLC into 72 fractions, which were recombined in a concatenated fashion into 6 sub-fractions. Each of the 6 sub-fractions was individually subjected to IMAC enrichment and lyophilized [36, 37]. Prior to LC-MS/MS analysis, all peptides were solubilized in 0.1% TFA containing iRT peptides (Biognosys AG, iRT).

***Mass spectrometry data acquisition.*** Samples were randomized and analyzed on an Exploris 480 mass spectrometer (Thermo) coupled with an Ultimate 3000 nano UPLC system and an EasySpray source. An estimated 2μg total peptide was analyzed for each whole proteome, and <2μg total phosphopeptide for each phosphoproteome sample. These were loaded onto an Acclaim PepMap 100 75um x 2cm trap column (Thermo) at 5μL/min, and separated by RP-HPLC on a nanocapillary column, 75 μm id × 50cm 2um PepMap RSLC C18 column (Thermo). Mobile phase A consisted of 0.1% formic acid and mobile phase B of 0.1% formic acid/acetonitrile. Peptides were eluted into the mass spectrometer at 300 nL/min, with each RP-LC run comprising a 90 min gradient from 3% B to 45% B. Data independent acquisition (DIA) mass spectrometer settings included one full MS scan at 120,000 resolution and a scan range of 350-1200 m/z with a normalized automatic gain control (AGC) target of 300% and an automatically determined maximum inject time. This was followed by variable (DIA) isolation windows, 30,000 resolution, a normalized AGC target of 1000%, and auto injection times. The default charge state was 3. The first mass was fixed at 250 m/z and the normalized collision energy for each window was 27.

We used data dependent acquisition settings for quality control analyses. For these control experiments, the mass spectrometer was set with a master scan at R=120000, a scan range of 300-1400, AGC target set to standard, maximum injection time set to auto, and dynamic exclusion set to 30 sec repeat 1. Charge states 2-5 were included. The top 15 data-dependent MS/MS scans were collected at R=45000, first mass set to 120, normalized AGC target at 300%, maximum injection time=auto, HCD NCE set to 30.

***QA/QC and system suitability.*** The suitability of the Exploris 480 instrument was monitored using QuiC software (Biognosys) for the analysis of the spiked-in iRT peptides. For quality control monitoring, standard digested *E. coli* protein stock was injected between samples (one injection after every four biological samples) and data were collected in data dependent acquisition (DDA) mode. These data were analyzed in MaxQuant [39] and the output was subsequently visualized using the PTXQC [39] package to track the quality of the instrumentation.

***Mass spectrometry raw data processing.*** DIA MS/MS raw files were processed in Spectronaut 16 (Biognosys AG) [40]. We used a reference human proteome containing 20,385 canonical and reviewed isoforms from Uniprot appended with the list of 245 common protein contaminants. Trypsin was specified as the enzyme, with up to two possible missed cleavages. Carbamidomethyl of cysteine was specified as a fixed modification and protein N-terminal acetylation and oxidation of methionine were considered as variable modifications. For the phosphoproteome data, we added phosphorylation of Serine, Threonine, and Tyrosine as variable modifications. A false discovery rate <1% was set for peptide and protein identification. All other search parameters were set to default values. The phosphopeptide spectral library was also generated in Spectronaut using default settings.
